# Robust integration of weakly anchored spatial multi-omics

**DOI:** 10.64898/2026.06.10.731246

**Authors:** Chuyao Wang, Yonghao Liu, Zhikang Wang, Pinli Sun, Zhi Li, Jieming Li, Xueting Wang, Ke Chen, Qi Zou, Daoliang Zhang, Zheqi Hu, Yixuan Du, Binzhi Qian, Xiaoyue Feng, Zhiyuan Yuan, Renchu Guan

**Affiliations:** Key Laboratory of Symbolic Computation and Knowledge Engineering of the Ministry of Education, College of Computer Science and Technology, Jilin University, Changchun, China; Institute of Science and Technology for Brain-Inspired Intelligence, Center for Medical Research and Innovation, Shanghai Pudong Hospital, Fudan University Pudong Medical Center, Fudan University, Shanghai, China; Center for Integrative Spatial-Omics Research, Fudan University, Shanghai, China; Department of Pathology, the Second Hospital of Jilin University, Jilin University, Changchun, China; National Clinical Research Center for Laboratory Medicine, Department of Laboratory Medicine, The First Hospital of China Medical University, Shenyang, China; Fudan University Shanghai Cancer Center, Department of Oncology, Shanghai Medical College, The Human Phenome Institute, Zhangjiang-Fudan International Innovation Center, Fudan University, Shanghai, China

## Abstract

Spatial multi-omics holds great promise for dissecting complex biological processes, though inherent technical constraints continue to limit its widespread adoption. Currently, most studies therefore measure distinct omics features on separate tissue sections, necessitating spatial diagonal integration. An emerging practical solution is to leverage hematoxylin and eosin (H&E) images as an integration anchor, given their ubiquity, low cost, and compatibility across tissue preparations. However, this anchor is frequently compromised in real-world settings by variations in H&E staining style, absence of reliable histological landmarks, and mismatches in spatial resolutions across omics modalities. To address this, we introduce SpaWeaver, a computational framework that couples a pathology foundation model with a graph Transformer and a latent feature aligner module, providing a highly robust solution for weakly anchored spatial omics data diagonal integration. Extensive experiments demonstrate that SpaWeaver exhibits superior robustness against isolated or synergistic weak-anchoring factors. The spatial multi-omics profiles generated by SpaWeaver link molecular features originally separated on two sections, unlocking diverse downstream analyses once exclusive to co-assayed spatial multi-omics data, including niche-aware cell–cell communication inference and multi-omics resolved cell state. In this study, it unveils tumor-distance-dependent fibroblast–CD4^+^ T-cell signaling in human colon adenocarcinoma and identifies a hypoxic glycolytic tumor state with pyknotic nuclei in human ovarian cancer. Overall, our approach bridges readily accessible single-omics measurements across weakly anchored tissue sections, enabling unified spatial multi-omics characterization and system-level tissue analysis.

## 1. Introduction

Biological processes are orchestrated across multiple molecular layers, such as chromatin accessibility, transcription, and protein abundance, with each omics modality offering distinct biological insights^1^. Emerging spatially resolved multi-omics sequencing technologies have been developed to profile complementary molecular modalities in the spatial context^2–4^, enabling the interrogation of regulatory and functional programs within complex tissues.

Performing multiple assays on the same tissue section typically necessitates delicate experimental workflows and stringent tissue preservation conditions. Critically, each new omics combination demands a bespoke experimental protocol, limiting flexibility and scalability. These challenges have spurred innovation along two complementary and mutually reinforcing tracks: continued experimental development of integrated spatial multi-omics assays, and computational strategies that reconstruct multi-omics landscapes from separately measured data^5–7^. Notably, spatial single-omics have already reached considerable maturity, offering high sensitivity, well-established protocols, and broad laboratory accessibility. Building on this foundation, a compelling computational paradigm has emerged: diagonal spatial omics integration, where adjacent tissue sections are profiled via distinct modalities and subsequently unified in silico^8–10^. This approach inherits the sensitivity and accessibility of mature single-omics platforms while remaining flexible enough to bridge any combination of modalities without a dedicated co-assay protocol. However, this strategy raises a fundamental challenge: how to reliably anchor and align omics features across sections that share no directly measured molecular features.

Computational integration has emerged as a key strategy to reconstruct a comprehensive multi-omics landscape from diagonally measured data. A straightforward strategy is to establish explicit links between molecular features across modalities using prior biological knowledge. Exemplifying this approach, SWITCH^11^ integrates spatial transcriptomics (ST) and chromatin accessibility by constructing peak-gene links based on genomic proximity^12^. The effectiveness of this strategy, however, is inherently constrained by the completeness and tissue-specificity of available prior knowledge, as the underlying regulatory landscape varies considerably across biological contexts and may not be fully captured by proximity-based rules alone. Recently, a growing body of work has demonstrated that unmeasured omics layers can be inferred directly from H&E images, with the learned associations being context-specific and data-driven rather than dependent on static cross-omics prior knowledge. This trajectory highlights histological features as a versatile and scalable anchor for spatial omics integration. Building on this momentum, SpatialEx+^9^ demonstrated the viability of extending this idea to spatial diagonal integration, using histological features as a common anchor to align omics measurements across distinct tissue sections.

Yet, relying on histology as a universal bridge inherently introduces unique vulnerabilities. In real-world settings, this histology anchor is frequently weakened by three practical factors: H&E style variation, missing histology anchors^13^, and mismatched spatial resolutions^14^. First, discrepancy in tissue pretreatment, dye composition, staining duration, scanner brightness, and imaging settings can conspire to induce severe domain shifts in H&E styles^15^, thereby rendering computational models highly susceptible to capturing these site-specific signatures. Second, many spatial omics platforms exhibit a bidirectional incompatibility with the standard H&E workflow. Specifically, the latter may induce biochemical alterations in tissue, compromising downstream molecular measurements; conversely, imaging-based spatial omics workflows may introduce chemical damage during repeated imaging cycles and mechanical damage due to the final removal of flow-cell^16^. Third, these challenges are further exacerbated by the inherent mismatch in spatial resolutions across spatial omics protocols (e.g., spot-level barcode-based ST^17^ versus single-cell-level spatial proteomics^18^). Together, we refer to settings in which one or more of these factors are present as **weakly anchored** spatial omics scenarios. These weakly anchored factors undermine the fundamental premise of H&E-anchored spatial omics generation methods, making existing computational models prone to capturing technical confounding rather than genuine biological variations.

In this study, we propose **SpaWeaver**, a unified framework for robust integration of weakly anchored spatial multi-omics data. SpaWeaver couples a pathology foundation model^19^ with a graph Transformer^20^ to encode cellular morphology together with the intrinsic spatial context, and incorporates a latent feature alignment strategy that integrates distinct tissue sections despite the severe domain shifts^21^ induced by weak anchoring. In a comprehensive benchmark on simulated datasets, SpaWeaver substantially surpassed the existing H&E-anchored spatial omics generation methods. Across four representative weakly anchored scenarios in real world, defined by two histology-anchor challenges (H&E style variation and missing histology anchors), crossed with matched and mismatched resolutions^22^, SpaWeaver achieved robust diagonal integration while preserving biological signals, facilitating downstream analysis. Leveraging the comprehensive multi-omics landscapes reconstructed by SpaWeaver, we identified tumor-distance-dependent communications between fibroblasts and CD4^+^ T cells in the human colon adenocarcinoma (COAD) microenvironment. In human ovarian cancer (OV) tissue, SpaWeaver uncovered a spatial domain enriched for an epithelial subpopulation characterized by hypoxic glycolytic programs and pyknotic nuclei, together with a lipid-metabolism-associated macrophages enriched domain that were frequently located nearby.

## 2. Results

### Overall architecture of SpaWeaver

SpaWeaver is designed for robust spatial diagonal integration (Fig. 1a) under diverse weakly anchored scenarios. In many spatial multi-omics studies, different molecular modalities are measured on adjacent tissue sections rather than on the same section, owing to the limited flexibility and availability of current co-assay technologies. From a computational perspective, this design produces a diagonal-like data structure: each section contains measurements for only a subset of molecular features. Spatial diagonal integration aims to complete this incomplete matrix by inferring the missing molecular profiles across sections, thereby generating *in silico* co-assayed spatial multi-omics data.

**Fig. 1:**
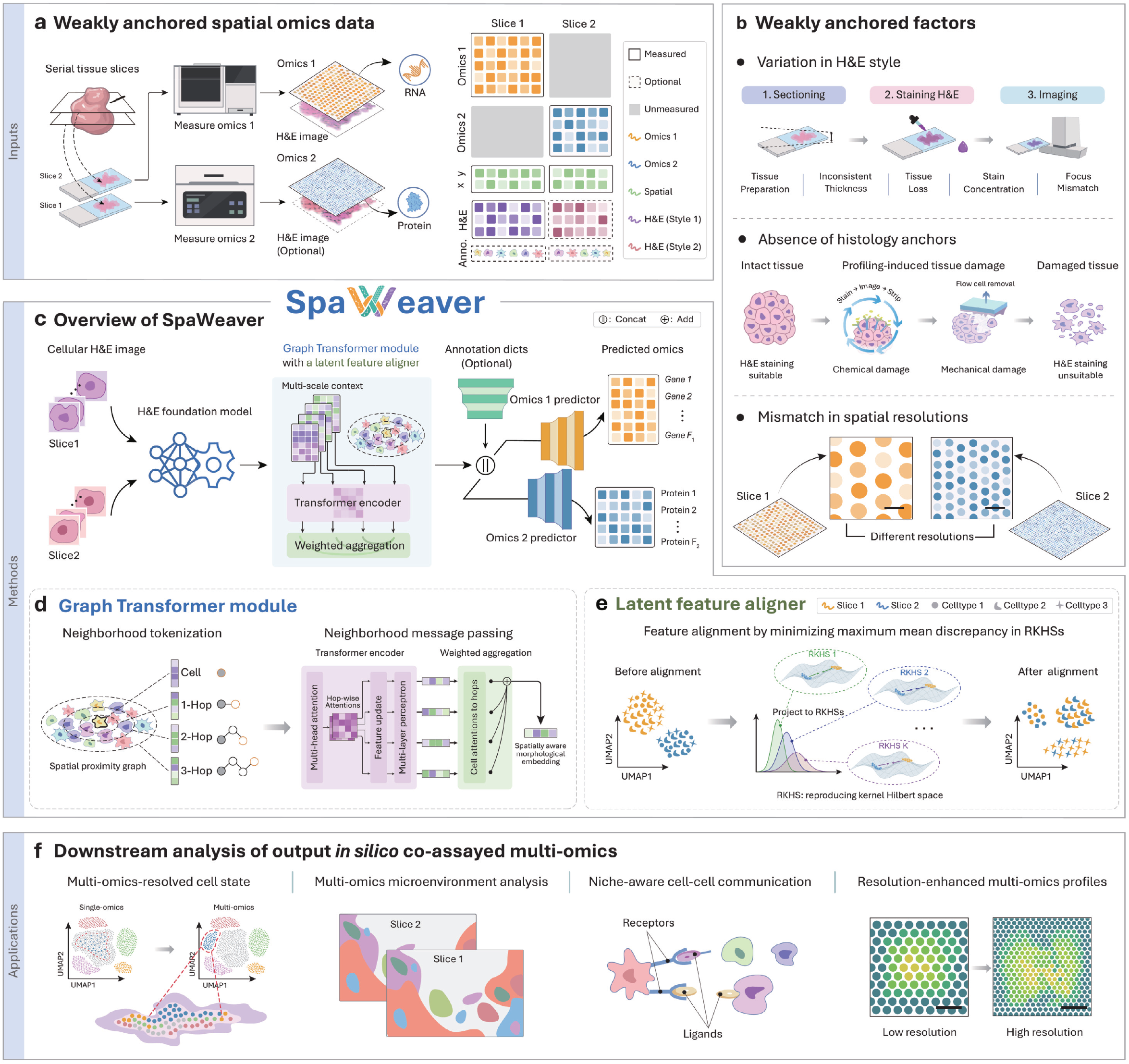
Overview of SpaWeaver workflow. **a**. Acquisition of weakly anchored spatial omics data. **b**. Three representative weakly anchored factors: H&E style variation, missing histology anchors, and mismatched spatial resolutions. **c**. Overview of the SpaWeaver framework. Cellular H&E patches are first embedded by a pretrained H&E foundation model, followed by a graph Transformer equipped with a latent feature aligner to derive cellular representations. These representations are subsequently combined with cell annotations (optional) and fed into two omics decoders to predict spatial omics profiles. **d**. Graph Transformer module, comprising cellular neighborhood tokenization and neighborhood message passing, enabling the encoding of cellular contexts at different scales. **e**. Latent feature aligner, which projects latent representations into RKHSs, then reduce domain shift via minimizing maximum mean discrepancy (MMD). **f**. The in silico co-profiled multi-omics data generated by SpaWeaver enable a range of downstream biological applications, including multi-omics-resolved cell state, multi-omics microenvironmental analysis, niche-aware cell–cell communication, and resolution-enhanced multi-omics profiles.

A common strategy is to use H&E images from adjacent sections as a shared anchor. Existing H&E-anchored spatial omics generation methods^23,24^ typically learn mappings from histological features to omics profiles within each section and then apply these mappings across sections to infer the unmeasured modalities. Some methods, such as SpatialEx+, further regularize the training of modality-specific predictors with additional alignment objectives. However, these strategies build on a reliable histology anchor across sections. In practice, this assumption is often weakened by H&E style variation, missing histology anchors, and mismatched spatial resolutions across spatial omics platforms (Fig. 1b). We refer to these settings as weakly anchored scenarios.

SpaWeaver addresses these challenges through a sequential architecture, it first extracts high-dimensional morphological representations from cell-, bin-, or spot-centered patches using a pre-trained H&E foundation model. These feature representations are then fed into a graph Transformer module equipped with a latent feature aligner (Fig. 1c). Conventional approaches typically rely on local message-passing mechanisms to capture neighborhood context^25,26^, but such strategies are constrained in modeling broader, tissue-level microenvironmental factors and long-range intercellular interactions that may shape cellular states beyond the immediate neighborhood^27^. To transcend these limitations, SpaWeaver adopts a graph Transformer architecture that enables multi-scale spatial context modeling (Fig. 1d, see Methods “Graph Transformer module”), conceptualizing cells analogously to tokens in natural language^28^. Specifically, a neighborhood tokenization strategy is employed to construct feature sequences for each cell, then neighborhood-aware message passing is subsequently leveraged to capture multi-scale spatial dependencies.

Next, to mitigate the domain shifts in weakly anchored data, these spatially aware histological representations are further regularized by minimizing the maximum mean discrepancy (MMD)^29^ between sections in reproducing kernel Hilbert space^30^ (Fig. 1e, see Methods “Latent feature aligner”). By doing so, cells with a similar biological context across sections are mapped close together in the feature space. To enhance biological specificity, auxiliary information such as cell type annotations can be incorporated as trainable embeddings at this stage, when available (see Methods, “Omics predictor”). Finally, modality-specific decoders translate the aligned representations into spatial omics profiles. The fully trained SpaWeaver enables spatial multi-omics feature inference across weakly anchored sections, facilitating downstream applications (Fig. 1f), including multi-omics-resolved cell state characterization, multi-omics microenvironment analysis, niche-aware cell–cell communication, and resolution-enhanced multi-omics profiles.

### Comprehensive benchmarking of panel diagonal integration under H&E style variation

To evaluate SpaWeaver’s resilience to H&E image style variation during the panel diagonal integration, we established a benchmark using a human breast cancer dataset profiled by Xenium^31^ (Fig. 2a, top). This dataset measured the same 313-gene panel on two serial sections, both paired with H&E staining, providing a well-controlled setting for quantitative evaluation. Specifically, we first split the measured gene panel into two non-overlapping subsets, assigning panel A (150 genes) to slice 1 and panel B (163 genes) to slice 2, while withholding the remaining genes in each slice to serve as the ground truth for quantitative evaluation (Fig. 2a, bottom). Originally, the two H&E images were generated in the same staining batch, they showed similar staining style and image quality (Supplementary Fig. 1). We then applied a staining perturbation algorithm^32^ on the H&E image from slice 2 (Fig. 2b, top) to simulate a weakly anchored scenario with H&E style variation. Uniform manifold approximation and projection (UMAP) visualization^33^, quantitative evaluation via scIB^34^ metrics, and the recently proposed scGraph^35^ (see Methods “Evaluation”) consistently demonstrated that this synthetic perturbation introduced a pronounced domain shift in cellular histological representations produced by the H&E foundation model (Fig. 2b, bottom).

**Fig. 2:**
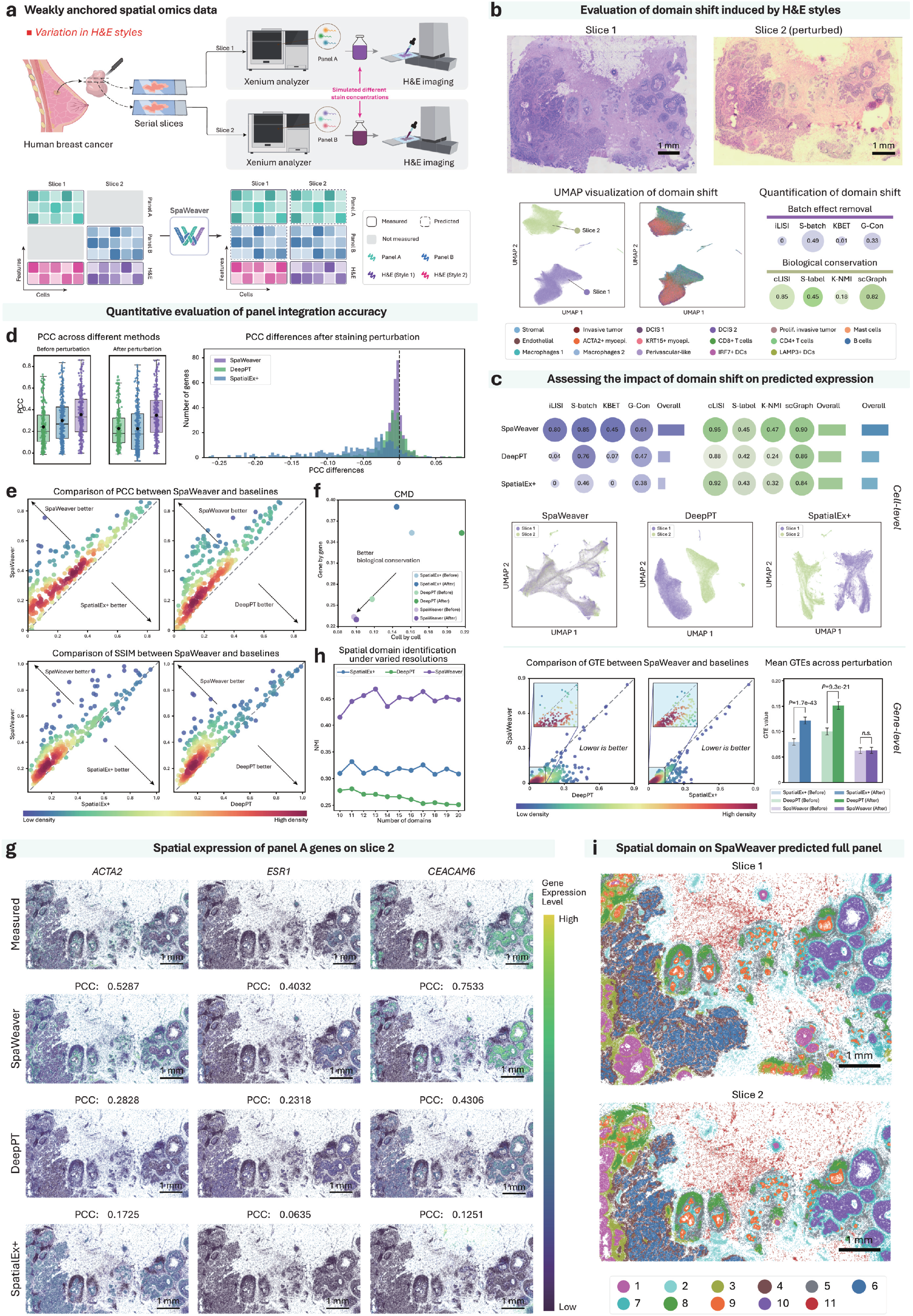
Benchmarking SpaWeaver panel diagonal integration under H&E style variation. **a**. Experimental design and computational workflow. Top: acquisition of the weakly anchored data with H&E style variation, where two consecutive breast cancer tissue sections were profiled with distinct, non-overlapping gene panels using Xenium, followed by subsequent H&E imaging subjected to varying stain concentrations. Bottom: the end-to-end computational flow of the SpaWeaver framework. **b**. Evaluation of domain shifts induced by H&E styles. Top: raw H&E image of slice 1, and staining perturbed H&E image of slice 2. Bottom left: UMAP visualization of the extracted cellular histology representations, colored by slice index (left) and cell type annotations (right). Bottom right: quantification of staining-induced domain shifts in the cellular histology representations. (prolif.: proliferating, myoepi.: myoepithelial) **c**. Benchmarking of the impact of domain shift on the gene expression predicted by each method. Top to bottom: quantification benchmarking; UMAP visualizations colored by slice index; scatter plots comparing gene-level GTE values between SpaWeaver and comparison methods colored by point density, with each point representing one of 313 genes; and bar plots showing the mean GTE values before and after the staining perturbation, with error bars representing the standard error of the mean (SEM). Two-sided paired Wilcoxon signed-rank tests were used, with *P* values shown (*n*=313 genes). **d**. Panel integration accuracy evaluation. PCC distributions before and after staining perturbation, along with the corresponding PCC differences. Box plots show median, mean, quartiles and 1.5× interquartile range. **e**. Scatter plots comparing PCC and SSIM values between SpaWeaver and baselines, with each point representing one of 313 genes, and colored by point density. **f**. Scatter plot of CMD values calculated across both gene and cell dimensions. **g**. Spatial expression of genes predicted by each method, with measured expression shown as reference. **h**. Similarity of spatial domains delineated between the measured and predicted panel B expression under varying domain numbers, quantified by NMI. **i**. Joint spatial domains on SpaWeaver predicted full panel expression. Parts of the figure were drawn by using pictures from Biovisart.

We first evaluated whether this stylistic domain shift distorted the predicted molecular profiles by comparing SpaWeaver with 8 methods supporting histology anchored spatial diagonal integration (SpatialEx+^9^, DeepPT^36^, iStar^37^, sCellST^24^, MISO^38^, HiST^39^, THItoGene^40^, and Hist2ST^41^). Quantitative comparison demonstrated that SpaWeaver consistently outperforms competing methods in both batch-effect removal and biological conservation (Fig. 2c, top, and Supplementary Table 1). Notably, regarding iLISI, most baseline methods showed values close to zero, whereas SpaWeaver reached 0.8. Regarding biological conservation, several baselines underperformed relative to the original cellular histological representations; in sharp contrast, SpaWeaver increased the K-NMI score by more than twofold. This indicates that our framework effectively attenuates staining-associated domain shifts without damaging intrinsic biological structure.

UMAP visualizations further corroborate these findings: expression profiles predicted by DeepPT and SpatialEx+ exhibited pronounced section-specific segregation, whereas SpaWeaver’s predictions were seamlessly blended (Fig. 2c, middle, and Supplementary Fig. 2a). We further quantified this shift at the gene level using gene transfer error (GTE)^42^, whereby lower values indicate smaller discrepancies between predicted and measured gene expression. SpaWeaver achieved a significantly lower GTE compared to DeepPT and SpatialEx+ (Fig. 2c, bottom, and Supplementary Fig. 2b). Collectively, SpaWeaver’s stable performance across severe staining variations underscores its superior robustness and practical utility.

Panel integration accuracy was quantitatively evaluated by comparing predicted and measured expression across the evaluated genes (Fig. 2d). Bar plots of gene-wise Pearson correlation coefficient (PCC) demonstrated that SpaWeaver achieves superior accuracy both before and after staining variation, while the histogram of PCC differences highlighted its robustness relative to DeepPT and SpatialEx+. A systematic comparison with the eight additional H&E-anchored prediction baselines further confirmed the consistent advantages of SpaWeaver (Supplementary Fig. 3). Furthermore, gene-by-gene comparisons utilizing both PCC and structural similarity index measure (SSIM) showed that SpaWeaver outperformed the baselines across most genes, indicating highly accurate imputation of both expression magnitude and spatial topological patterns (Fig. 2e, and Supplementary Fig. 4a). To evaluate whether the predicted gene expressions retained true biological relevance, we calculated correlation matrix distance (CMD) scores for all methods. Remarkably, SpaWeaver yielded the lowest CMD values from both gene and cell perspectives (Fig. 2f, and Supplementary Fig. 4b), validating its capacity to preserve intrinsic gene co-expression networks.

Furthermore, we visualized the spatial expressions of the three representative genes predicted on slice 2, including *ACTA2, ESR1*, and *CEACAM6* (Fig. 2g and Supplementary Fig. 5). Notably, SpatialEx+ accurately predicted *ACTA2* (a canonical marker for cancer-associated fibroblasts, or CAFs), before staining shift, but erroneously imputed aberrant *ACTA2* expression within tumor epithelial regions after staining shift. This pivotal observation reveals that H&E style variation does not merely induce a conventional batch effect; rather, they can catastrophically corrupt the learned histology-to-omics mapping through representation-level semantic misalignment, thereby distorting spatial gene-expression patterns. Similarly, *CEACAM6* is endogenously detected in ductal carcinoma in situ (DCIS) regions, but DeepPT predicts prominent ectopic signals within invasive ductal carcinoma (IDC) regions. In contrast, SpaWeaver successfully recovered these spatially restricted patterns. Remarkably, SpaWeaver remained highly effective even for *ESR1*, a weakly expressed but clinically critical hormone receptor associated gene for breast cancer molecular classification^43^. Specifically, SpaWeaver achieved a high PCC of 0.4032, substantially outperforming DeepPT (0.2318) and SpatialEx+ (0.0635).

To evaluate downstream biological utility, we conducted unsupervised spatial domain identification^44^ using the predicted expression profiles. Quantitative evaluation based on normalized mutual information (NMI) demonstrated that SpaWeaver consistently produced domain assignments that closely matched those derived from the experimental ground truth across a wide range of spatial resolutions (Fig. 2h). Finally, when performing spatial domain identification on the fully complemented panels predicted by each method, SpaWeaver produced consistent spatial domains across the two sections without observable batch effects (Fig. 2i). In contrast, DeepPT, SpatialEx+ and other comparison methods yielded discordant domain delineations across slices due to staining variation (Supplementary Fig. 6). We further applied scVI batch correction^45^ to the gene expression profiles predicted by the comparison methods, but this post hoc correction failed to resolve the domain shifts induced by H&E style variation. These results suggest that the impact of H&E style variation is not a simple downstream batch effect in the predicted expression space, but instead reflects distortions introduced during histology-anchored omics inference.

In this experimental setting, we also validated the individual contribution of each constituent module within SpaWeaver, as detailed in Supplementary Fig. 7. Taken together, our benchmarks demonstrated that staining-style variations, which are ubiquitously encountered in real-world multi-section studies, can induce severe semantic misalignment in panel diagonal integration. Crucially, this vulnerability may compromise other conventional H&E-anchored spatial omics generation pipelines. By effectively mitigating this misalignment, SpaWeaver maintained highly accurate and robust spatial omics prediction even under such severely perturbed, weakly anchored scenarios.

### Cross-resolution panel diagonal integration in weakly anchored spatial omics data

A more stringent weakly anchored scenario was then considered, where H&E style variations and mismatched spatial resolutions exist simultaneously (Fig. 3a, top). This benchmark used a serial human colorectal cancer (CRC) sections, with slice 1 measured by Xenium and slice 2 measured by Visium HD (8 *μ*m bin used)^46^. The shared 402 genes between the two slices were divided into two non-overlapping panels, where panel A was assigned to slice 1, and panel B was assigned to slice 2. This setting mimics a diagonal integration scenario with mismatched spatial resolutions, while the withheld genes from training on each slice can be used as reference to evaluate prediction accuracy (Fig. 3a, bottom). Together with the inherent H&E style variation between the two sections which is likely stemming from platform-specific sample preprocessing and institutional differences in staining protocols (Fig. 3b), this benchmark constituted a more challenging weakly anchored scenario. Consistently, UMAP visualization of the histological representations demonstrated a pronounced domain shift between the two sections.

**Fig. 3:**
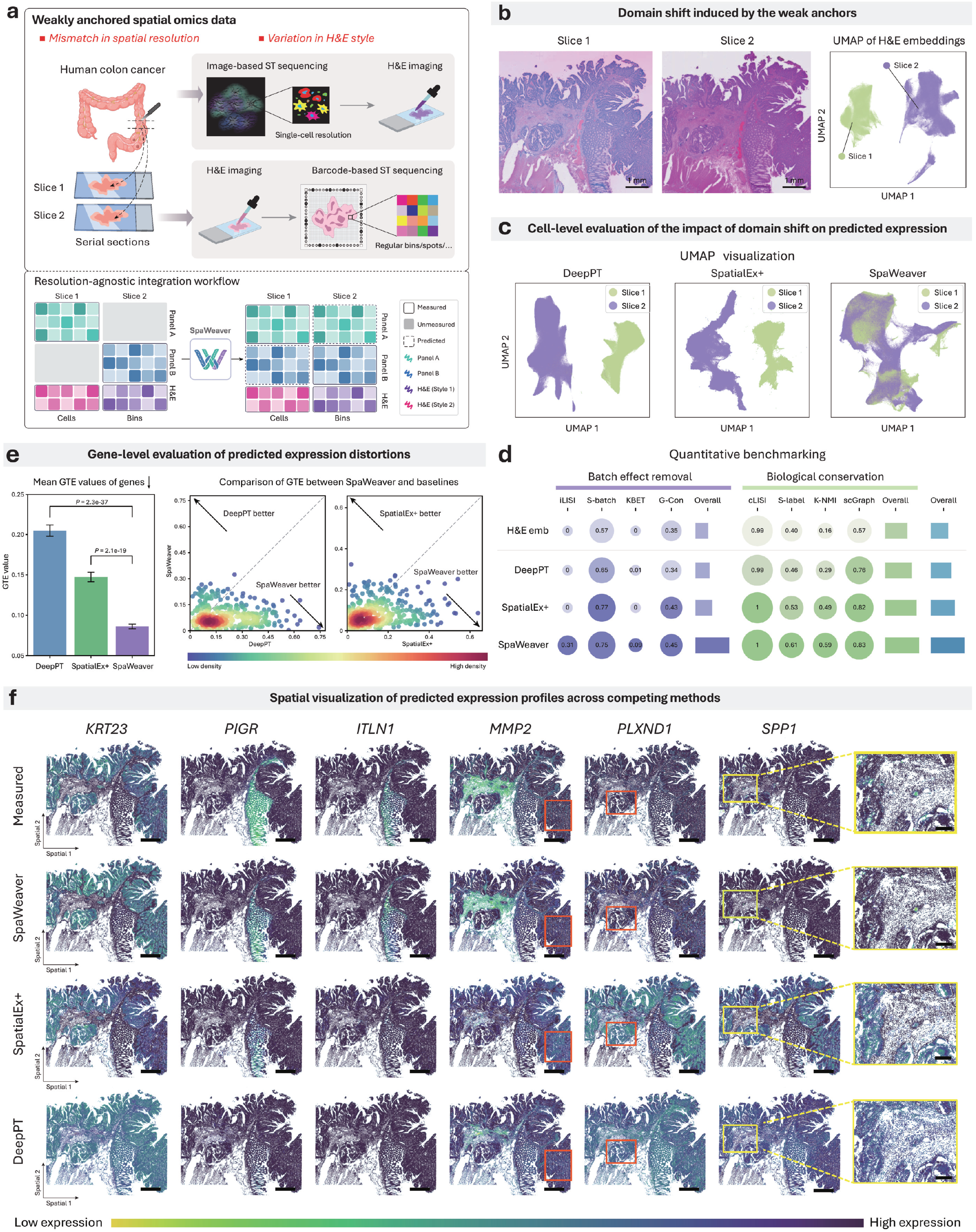
SpaWeaver panel diagonal integration with mismatched resolutions. **a**. Top: acquisition of the weakly anchored data with H&E style variation and mismatch spatial resolutions, where two serial human CRC tissue sections were profiled using different ST protocols. Bottom: the end-to-end computational workflow of SpaWeaver. **b**. H&E images on the two slices, and UMAP of cellular image representations colored by slice identity. **c**. UMAPs of gene expression predicted by each method, colored by slice index. **d**. Quantitative benchmarking the impact of weak anchors on the predicted gene expression from each method. **e**. Left: Bar plots of mean GTE values for different methods, with error bars representing the SEM. Two-sided paired Wilcoxon signed-rank tests were used, with *P* values shown (*n*=402 genes). Right, scatter plots of GTE values between SpaWeaver and comparison methods, with each point represents one of 402 genes, and colored by point density. **f**. Spatial visualization of gene expression predicted from each method on slice 2, with measured expression shown as reference. Scale bars, 1 mm (**f**) and 150 *μ*m (zoomed-in **f**).

DeepPT and SpatialEx+ were selected as the representative baselines for this benchmark due to their competitive performance in the previous evaluation setting. We then assessed the cross-section consistency between the predicted profiles on the two adjacent tissues. UMAP visualization revealed that the predictions from DeepPT and SpatialEx+ exhibited persistent, slice-specific segregation, indicating that the baseline histology-to-omics mappings remained heavily distorted by the technical domain shift (Fig. 3c). Quantitative evaluation utilizing both data integration and biological conservation metrics further supported this observation: SpaWeaver achieved the most robust balance between mitigating section-specific batch effects and preserving biological fidelity (Fig. 3d). Furthermore, SpaWeaver yielded the lowest mean GTE, significantly outperforming DeepPT and SpatialEx+. Pairwise comparison of individual GTE values demonstrated that most genes were below the identity diagonal when SpaWeaver was benchmarked against DeepPT or SpatialEx+ (Fig. 3e). This pivotal finding confirmed a global, gene-wise superiority rather than an artifactual improvement driven by a small subset of highly variable transcripts.

Spatial visualization of representative withheld genes revealed distinct failure modes among the baseline methods (Fig. 3f). Taking *KRT23* as an example, DeepPT produced weakly structured predictions with reduced spatial contrast. This imputation deficit could be attributed to its lack of explicit spatial-context modeling, which fundamentally limits its ability to leverage microenvironmental information to infer or reinforce molecular signals under sparse omics measurements, such as the 8 *μ*m Visium HD data utilized here. In contrast, SpatialEx+ often generated spatially organized expression patterns, for example *PIGR*, consistent with its relatively strong biological conservation scores in the quantitative evaluation. However, these predicted patterns were frequently assigned to incorrect tissue regions, underscoring a severe semantic misalignment under weakly anchored settings. Remarkably, SpaWeaver successfully bypassed both limitations, delivering predictions that were both highly structured and faithfully aligned with the experimentally measured expression patterns. Beyond genes with distinct, continuous spatial layout, SpaWeaver also remained robust in more challenging scenarios. These included *PLXND1*, which exhibited low expression intensity overall, and *SPP1*, whose expression was restricted to a small niche of macrophages but is clinically linked to reduced CD8^+^ T cell cytotoxic activity in CRC^46^.

### Challenge of spatial multi-omics diagonal integration without histology anchor

In spatial multi-omics studies, proteomics provides essential protein-level information that enables a more direct characterization of cell identities, functional states, and tissue microenvironment organization beyond transcriptomics alone. For example, tumor immunology studies often profile ST on one tissue section and spatial proteomics on an adjacent section to jointly characterize immune cell states and tumor-immune interactions. However, spatial proteomics datasets are commonly generated without a matched H&E image, because antibody-based staining, iterative imaging-stripping workflows are optimized for protein detection and may compromise tissue morphology (Fig. 4a). This phenomenon is widespread across representative benchmarks from SPATCH^47^ and STP-Portal^13^, spanning diverse tissue types and multiplexed spatial proteomics platforms such as CODEX^18^ and IBEX^48^ (Fig. 4b). Although adjacent transcriptomic and proteomic sections provide a nominal pair, the absence of a shared histology anchor, compounded by non-overlapping molecular features, makes this correspondence intrinsically weak. This challenging diagonal integration setting remains beyond the scope of existing methods.

**Fig. 4:**
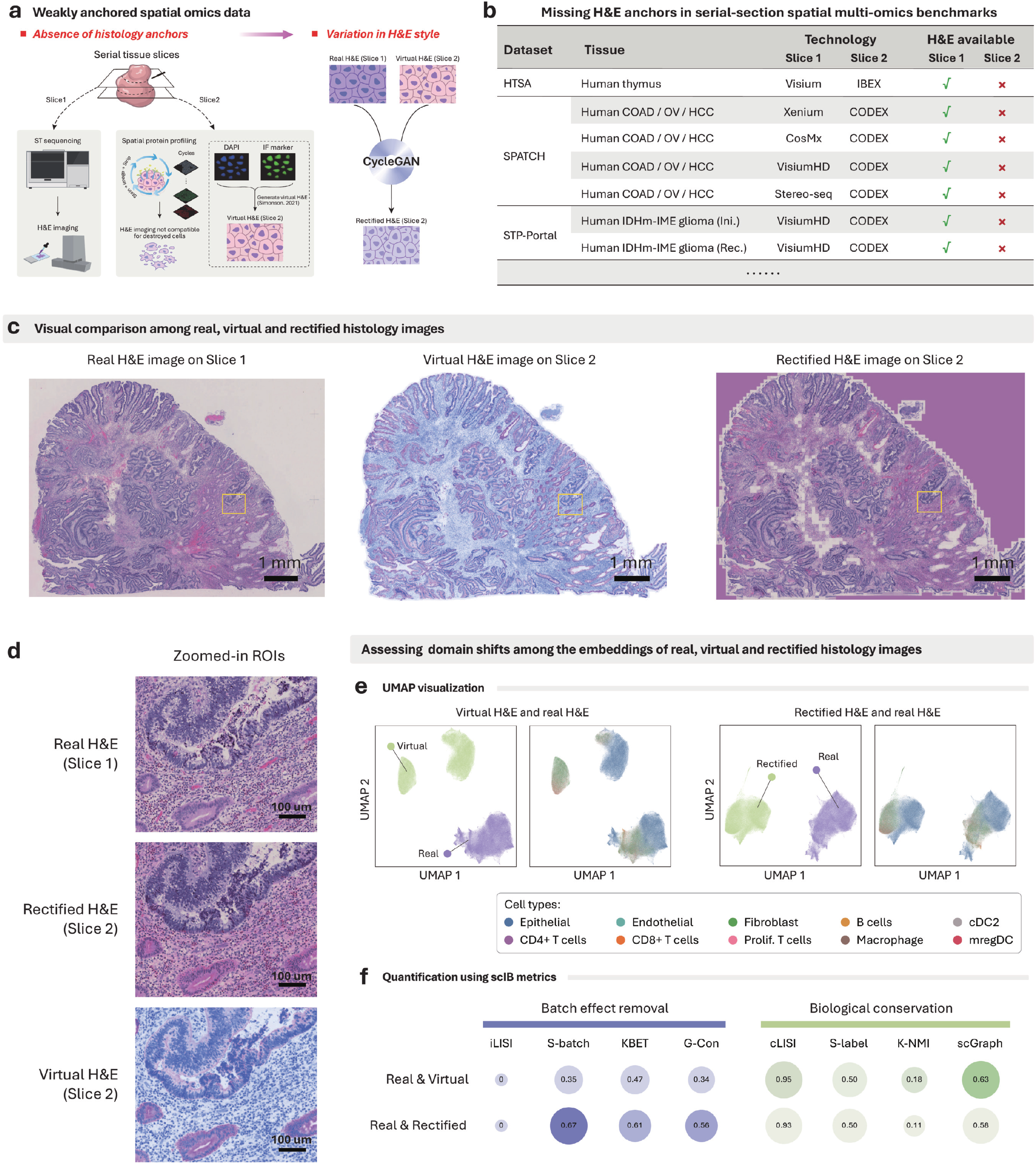
Challenges of multi-omics diagonal integration without histology anchor. **a**. Schematic workflow transforming the missing histology anchor scenario into an H&E style variation problem via virtual H&E image generation. **b**. Survey of public spatial multi-omics datasets highlighting the pervasive absence of co-stained H&E images. (Ini.: initial; Rec.: recurrent) **c**. Visual comparison among real, virtual, and rectified H&E images. **d**. Zoomed-in ROIs (Yellow boxes in **c**) of real, rectified and virtual H&E image, respectively. **e**. UMAPs of cellular morphology representations, colored by slice index and cell type annotations. (prolif.: proliferating) **f**. Quantitative comparison of domain shift in morphology representations extracted across real, virtual, and rectified H&E images.

Notably, the multiplexed spatial proteomics channels contain native, morphology-related signals that inherently resemble key components of classical H&E staining. Specifically, DAPI nuclear staining, routinely acquired for single-cell segmentation, captured explicit nuclear morphology, while cytoplasmic and membrane protein markers outline coarse cellular contours. These intrinsic biological signals can therefore be leveraged to synthesize virtual H&E images^14^. As an illustrative example, we utilized a COAD tissue section profiled by CODEX from the SPATCH dataset to generate virtual H&E images; however, the raw synthetic outputs showed substantial style variations compared with the real H&E images acquired from the adjacent slice (Fig. 4c, center). Given that image style transfer is well-established in computer vision and CycleGAN^49^ has been widely used for unsupervised image-to-image translation without requiring paired training data, we applied CycleGAN to align the virtual H&E images into the exact visual style of the real H&E, a process we referred to as virtual H&E rectification, yielding “rectified H&E” (Fig. 4c, right). Remarkably, the resulting rectified H&E images closely resembled the true experimental H&E references upon visual inspection, preserving both macroscopic tissue architecture and finer morphological details (Fig. 4d).

We next extracted cellular histological representations by passing cellular patches from the real, virtual, and rectified H&E images through the H&E foundation model, enabling a direct comparison of their feature spaces before and after style rectification (Fig. 4e). Strikingly, UMAP visualization revealed that despite visual similarities, the cellular representations from both the virtual and rectified H&E images exhibit a pronounced, systematic separation from those generated by the real H&E patches (Fig. 4e). These empirical findings demonstrated that while CycleGAN succeeded in optimizing macro-level style stylization, it introduced subtle, watermark-like artifacts that, although imperceptible to human visual inspection, fundamentally contaminated the deep embedding space of the H&E foundation model, leading to a severe domain shift. We systematically quantified the magnitude of these artifacts utilizing scIB benchmarking metrics and scGraph (Fig. 4f). Although style rectification improved select batch-effect removal metrics (such as S-batch, KBET and G-Con), alternative alignment scores remained near zero, confirming the persistence of the underlying domain shift. Concurrently, the rectified H&E exhibited a degradation across all biological conservation metrics, with k-NMI scores collapsing by seven percentage points.

Collectively, these findings demonstrated that CycleGAN-based style rectification largely fails to resolve the representation-level domain shift between virtual and real H&E images, and can even inadvertently disrupt biologically meaningful structure. However, the relatively favorable biological conservation scores between the real and virtual H&E image representations (Fig. 4f) strongly suggested that the unrectified virtual H&E still retains crucial, high-fidelity morphological information. This intrinsic structural integrity opens a new avenue, providing a vital structural context to anchor multi-omics diagonal integration under a scenario completely devoid of experimental histology anchor.

### Spatial multi-omics diagonal integration without histology anchor

Given that spatial proteomics datasets commonly lack co-stained H&E images, we next considered a weakly anchored scenario without histology anchor (Fig. 5a). The same COAD dataset in the previous section was adopted for this benchmark. Specifically, the dataset comprises two serial slices, with slice 1 being profiled by Xenium and paired with a co-stained H&E image, whereas slice 2 being profiled by CODEX but lacked a co-stained H&E image. Building upon the insights in the previous section, SpaWeaver generated a virtual H&E image for slice 2 as the surrogate anchor. During model optimization, SpaWeaver flexibly incorporated prior information when available, and the readily accessible cell type annotations were utilized in this scenario (Fig. 5b). For the ST slice, these annotations can be derived using established ST annotation methods, such as Spatial-ID^50^. Conversely, considering that spatial proteomics panels typically profile only dozens of markers, we further developed a dedicated marker-based annotation strategy tailored specifically for spatial proteomics data (see Methods “Data”). The resulting annotations for slice 2 are detailed in Supplementary Fig. 8.

**Fig. 5:**
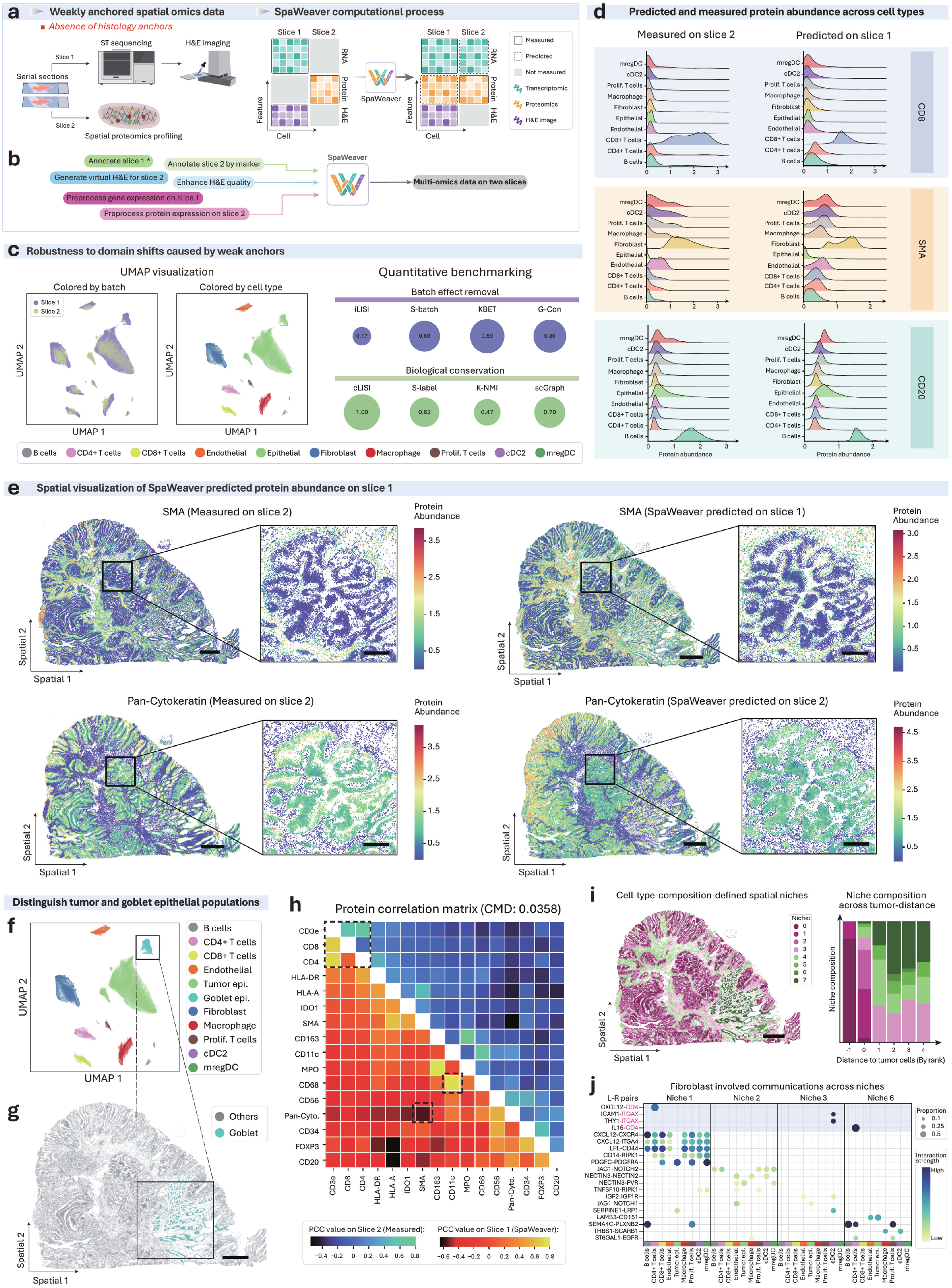
SpaWeaver multi-omics diagonal integration without histology anchor. **a**. Left: acquisition of the weakly anchored data without histology anchors, where slice 1 subjected to Xenium, with co-stained H&E image, and slice 2 only subject to CODEX. Right: The end-to-end computational workflow of SpaWeaver. **b**. Preprocessing of weakly anchored data prior to model training in SpaWeaver. **c**. Cross-slice concordance of multi-omics profiles visualized by UMAP (left) and quantification metrics (right). **d**. Ridge plots of SpaWeaver-predicted protein abundance across cell types on slice 1, with the distribution of measured proteomic on slice 2 shown as reference. **e**. Spatial visualization of SpaWeaver predicted abundance of SMA and Pan-Cytokeratin on slice 1, with the measured counterparts on slice 2 shown as reference. **f**. Multi-omics UMAP embeddings colored with new cell type. **g**. Spatial distribution of goblet cells. **h**. Heatmap of PCC values between the proteins computed from profiles predicted by SpaWeaver on slice 1 (bottom-left) and measured on slice 2 (top-right). **i**. Left: spatial niche delineation on tissue. Right: Niches composition across different distance rank to tumor cells. **j**. Fibroblast involved communications across distinct spatial niches, wherein ligand or receptor molecules highlighted in pink were imputed by SpaWeaver. Scale bars, 1 mm (**e, g, i**) and 250 *μ*m (zoomed-in **e**). (prolif.: proliferating; epi: epithelial)

We first generated UMAP visualization of the predicted multi-omics expression profiles of the weakly anchored slices (Fig. 5c). In contrast to the pronounced separation observed in Fig. 4e, SpaWeaver achieved effective integration of the two distinct slices while maintaining a clear, coherent cell-type organization. Quantitative evaluations further substantiated this observation: the KBET score reached 0.89, indicating strong batch mixing, accompanied by a scGraph score of 0.70, indicating high preservation of the underlying biological structures. We next compared the distributions of predicted protein expression on slice 1 with the experimentally measured counterparts on slice 2 across diverse cell types (Fig. 5d). Three representative proteins, including CD8, CD20, and SMA, were selected for detailed evaluation. Notably, SpaWeaver accurately captured cell-type-specific protein expression patterns, for example, CD8 was highly expressed in CD8^+^ T cells, CD20 was enriched in B cells, and SMA showed elevated expression in both fibroblasts and endothelial cells. Qualitative spatial distributions of SMA and Pan-Cytokeratin are presented in Fig. 5e. These spatial profiles showed that predictions by SpaWeaver closely recapitulated the experimental ground truth, with the high-magnification views of selected regions of interest (ROIs) further confirming their spatial concordance at a finer resolution. Additional benchmarking examples are provided in Supplementary Fig. 9a.

Notably, the multi-omics UMAP embedding revealed two distinct epithelial subpopulations. Utilizing K-Means clustering, we partitioned one of these subpopulations (Fig. 5f), whose corresponding spatial distribution is illustrated in Fig. 5g. Based on histological morphology and characteristic gene expression patterns, this cluster was annotated as goblet cells. Differential expression analysis further confirmed that this specific population significantly upregulated canonical secretory genes, including *FCGBP* and *DUOX2*^51^, whereas key tumor-associated genes, such as *IGF2*^52^, *TP53*^53^, *PCNA*^54^, and *FASN*^55^, were predominantly enriched in the remaining epithelial population (Supplementary Fig. 9b).

Finally, we leveraged the experimentally measured transcriptomics in conjunction with the predicted proteomics to perform cell–cell communication analysis on slice 1. We first evaluated the preservation of protein-protein correlation patterns between the predicted and measured proteomic profiles. As illustrated in Fig. 5h, SpaWeaver accurately preserved these co-functional topological structures, yielding a CMD of 0.0358. For instance, CD3e expression was positively correlated with both CD4 and CD8, whereas CD4 and CD8 exhibited minimal correlation with each other. This coordinated pattern precisely recapitulated the hierarchical marker structure of T cells: CD3e marks the pan-T-cell lineage, while CD4 and CD8 characterize largely mutually exclusive T cell subsets^56^. Moreover, the positive correlation between CD68 and CD11c was successfully preserved, reflecting their well-documented co-expression in myeloid-derived antigen-presenting cells, including macrophages and dendritic cells^57,58^. These validation outcomes confirmed that the imputed proteomic profiles successfully retained the native protein expression network, establishing a reliable cornerstone for downstream ligand-receptor interaction modeling.

To characterize local cell–cell communication patterns, we defined spatial niche based on neighborhood cellular composition (Fig. 5i, left), with the spatial distribution of these niches relative to the tumor epithelial cells illustrated in Fig. 5i, right. We then applied CellphoneDB^59^ on slice 1 to identify enriched ligand-receptor interactions (Fig. 5j), wherein specific ligand or receptor molecules highlighted in pink were imputed by SpaWeaver, while the remaining genes were experimentally measured. Taking fibroblasts as an illustrative example, we observed distinct niche-specific communication with CD4^+^ T cells. In niche 1, which is located adjacent to the tumor region, fibroblast-CD4^+^ T-cell interactions were primarily mediated by the CXCL12-CD4 signaling axis. In contrast, within niche 6, which is distal to the tumor region, IL16-CD4 signaling became dominant. Mechanistically, CXCL12 and IL-16 represent distinct routes of CD4^+^ T-cell recruitment. CXCL12 is a stromal chemokine that acts through the CXCR4 axis to regulate immune cell trafficking, a process frequently associated with immunosuppression^60^. By contrast, IL-16 directly binds to CD4 and primarily mediates the migration and activation of CD4^+^ T cells^61^. Together, these results demonstrated that the co-profiled transcriptomics and proteomics data derived by SpaWeaver provide a more comprehensive molecular context for resolving spatially organized cell–cell communication.

### Cross-resolution multi-omics diagonal integration without histology anchor

Having demonstrated SpaWeaver’s robustness in weakly anchored scenario without histology anchor, we next assessed its performance when this challenge was compounded by spatial resolution mismatch. SpaWeaver was applied to a stage IVB OV dataset comprising two adjacent sections, with ST measured by Xenium together with co-registered H&E imaging in slice 1 and spatial proteomics profiled by CODEX in slice 2 (Fig. 6a and Supplementary Fig. 10a)^47^. To simulate a cross-resolution setting, Xenium data were down-sampled to 55 *μ*m spots to approximate Visium resolution. To facilitate integration, SpaWeaver incorporated multiple sources of information, including cell-type proportion in slice 1 (the simulated Visium data) (Supplementary Fig. 10b) and coarse-grained cell-type annotations in slice 2 (Supplementary Fig. 10c). Following SpaWeaver integration, a resolution enhancement step was applied to reconstruct multi-omics profiles at single-cell resolution.

**Fig. 6:**
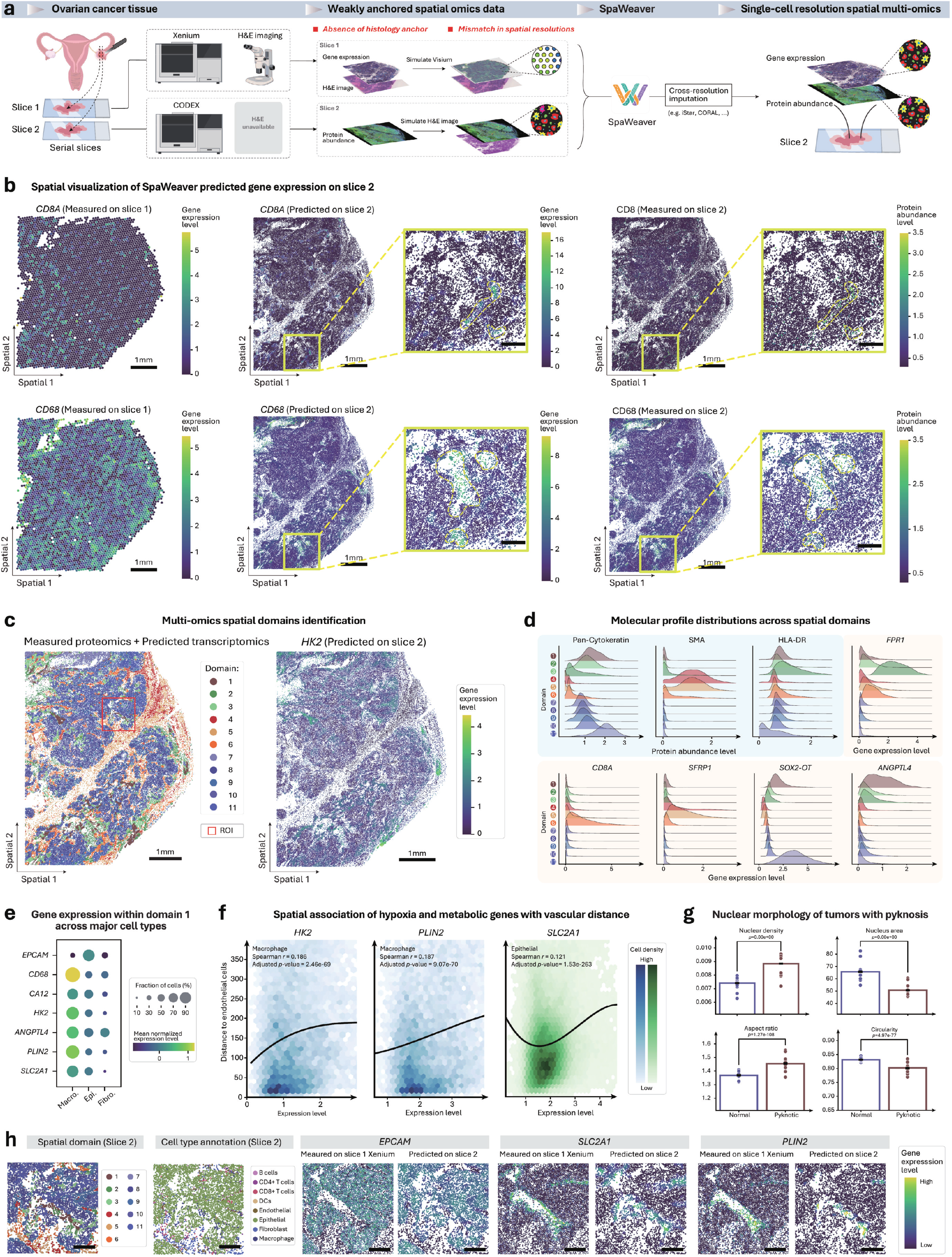
SpaWeaver cross-resolution multi-omics diagonal integration without histology anchor. **a**. From left to right: simulation of weakly anchored data with mismatch spatial resolutions from serial OV slices without histology anchors; These data put through SpaWeaver and super-resolution method can generate single-cell co-assayed multi-omics data in silico. **b**. Spatial visualization of predicted single-cell-resolution expression of gene *CD8A* and *CD68* on slice 2, with their coding proteins abundance measured on slice 2 shown as reference. **c**. Left: spatial domain identification using multi-omics embedding. Right: spatial expression of representative marker *HK2* for domain 1. **d**. Ridge plot of molecule feature distribution across spatial domains. **e**. Marker gene expression on major cell types within domain 1. (epi.: epithelial; fibro.: fibroblast) **f**. Hex-bin plots illustrating spatial association between the expression of hypoxia-associated metabolic genes and physical distance to the closest endothelial cell, overlaid with third-order polynomial fits and density-coded distributions. Statistical significance was evaluated via Benjamini–Hochberg-corrected Spearman rank correlation tests (adjusted *P* value and Spearman *r* explicitly indicated). **g**. Bar plots quantifying nuclear morphology differences between tumor cells in domain 1 and other domains, with error bars representing the SEM. Two-sided unpaired Wilcoxon rank sum tests were used to compare the differential, with *P* values shown. **h**. ROI of domains 1 and 2. From left to right: spatial domain distribution, cell type annotation, and imputed single-cell gene expression, with expression on slice 1 measured by Xenium shown as references.

We first compared SpaWeaver-predicted gene expression at single-cell resolution with the corresponding protein abundance measured on slice 2 (Fig. 6b). Canonical markers such as *CD8A* (CD8^+^ T cells) and *CD68* (macrophages) appear spatially dispersed without clear spatial organization on slice 1 at spot resolution, whereas SpaWeaver resolves fine-grained expression patterns consistent with their translated proteins (Supplementary Fig. 11a).

We next performed spatial domain identification using the measured protein abundance together with SpaWeaver-predicted transcriptomic profiles (Fig. 6c). Specifically, domains 4 and 5 were identified as stromal-associated regions, as indicated by elevated SMA abundance, whereas domains 3 and 6 represented immune-associated regions, indicated by high HLA-DR abundance. The remaining domains likely corresponded to tumor cores or tumor margins, as suggested by broad Pan-Cytokeratin abundance (Fig. 6d).

Notably, domain 1 could not be resolved by the measured proteomics alone, as no measured proteins exhibited a consistent spatial pattern (Supplementary Fig. 11b, c). This indicates that SpaWeaver-predicted transcriptomic data played a key role in defining this domain. Particularly, domain 1 is characterized by elevated expression of *CA12, HK2, ANGPTL4, PLIN2*, and *SLC2A1* (Supplementary Fig. 11d, e), with expression predominantly arising from tumor epithelial cells and macrophages (Fig. 6e). Functionally, these genes reflect key aspects of metabolic adaptation to hypoxia, including glycolysis (*SLC2A1, HK2*)^62^, pH regulation (*CA12*)^63^, and lipid metabolism (*PLIN2*)^64^. Using distance to endothelial cells as a proxy for vascular proximity, expression of these genes increased with increasing distance from vasculature (Fig. 6f), consistent with increased hypoxia stress in regions distal to blood supply. To further characterize domain 1, we selected 10 ROIs enriched for domain 1, and 10 tumor ROIs lacking domain 1 in slice 2 (Supplementary Fig. 11f), then registered these ROIs to the H&E image of slice 1 for morphological comparison. Tumor epithelial cells within domain 1 enriched ROIs exhibited increased nuclear density, reduced nuclear size, higher aspect ratio, and irregular nuclear morphology relative to other tumor regions (Fig. 6g). These features collectively indicated a subpopulation of tumor cells under hypoxic stress with elevated glycolytic metabolism^65^ and a pyknotic nuclear phenotype.

A representative ROI enriched for domain 1 is shown in Fig. 6h. Notably, domain 2, a pro-inflammatory (M1-like) macrophage-enriched domain, often appeared adjacent to domain 1 (Supplementary Fig. 11c, g) with similarly high expression of *PLIN2* and *SLC2A1*. We therefore speculated that a subset of tumor cells within domain 1 undergo necrosis under these extreme survival conditions, and the released lipids may attract infiltrating macrophages, thereby forming this spatial domain. While the temporal progression from pyknotic nuclei to cell death cannot be directly inferred from this static data, one possibility is that the tumor cells failing to withstand the combined stress of high proliferative density and hypoxia contribute to necrosis.

Overall, these results highlight the ability of SpaWeaver to integrate spatial multi-omics data despite the absence of H&E image anchor and the resolution differences across spatial omics protocols, thereby facilitating downstream multi-omics biological interpretation and discovery at high spatial resolution.

## 3. Discussion

A foundational objective of spatial multi-omics is to realize a unified, multi-dimension molecular characterization of complex tissues by concurrently resolving disparate omics layers within their native architectural contexts. However, despite the exponential diversification of spatial profiling technologies, the simultaneous experimental measurement of distinct molecular modalities on a single tissue section remains technically prohibitive, financially taxing, and inherently difficult to scale. Consequently, a vast majority of translational spatial studies partition different molecular layers across adjacent tissue sections, creating a critical open challenge for spatial diagonal integration.

Existing computational approaches often heavily rely on identical biological prior or shared H&E histology anchors to guide this integration. In real-world multi-omics experiments, however, this strict assumption is frequently violated: H&E images may exhibit substantial style variability, histology anchors may be entirely missing for certain modalities, and spatial resolutions may differ markedly across diverse omics assays. To render spatial diagonal integration broadly applicable in such realistic experimental settings, a dedicated framework capable of explicitly accommodating these weakly anchored spatial omics data is critically required. To bridge this technological chasm, we developed SpaWeaver, an AI-driven computational framework for the robust integration of weakly anchored spatial multi-omics. In rigorous benchmarking simulations mimicking severe H&E style variation, SpaWeaver outperformed eight state-of-the-art alternative algorithms by a substantial margin. Furthermore, across extensive evaluations in more complex, real-world scenarios, SpaWeaver maintained exceptional robustness even when multiple weak-anchoring challenges were intertwined.

However, several limitations of the current framework warrant acknowledgment. First, due to the practical bottleneck in acquiring benchmarks with three or more omics layers, SpaWeaver was not explicitly optimized for multi-section diagonal integration involving arbitrary tissue slices. We anticipate that the emergence of such multi-layered datasets will naturally catalyze the next generation of computational methods for these unconstrained scenarios, ultimately enabling a more holistic, pan-tissue-scale multi-omics synthesis. Second, although our evaluation revealed that image-generating models like CycleGAN may introduce subtle computational domain shifts, cross-modal image translation remains a promising avenue for future exploration, particularly for imaging-based spatial assays.

Despite these open avenues for optimization, SpaWeaver substantially broadens the computational horizons of spatial diagonal integration and democratizes the utility of experimentally accessible, single-modality spatial omics datasets. By reconstructing high-resolution spatial multi-omics within a single tissue slice in silico, SpaWeaver establishes a rich spatial molecular context for diverse downstream mechanistic discoveries. We envision that this framework will serve as a cornerstone for future integrative tissue atlases, fundamentally empowering comprehensive systems-biology exploration.

## 5. Methods

### The SpaWeaver model

SpaWeaver is a novel graph-based framework designed for the robust integration of weakly anchored spatial omics data. We denoted the molecular feature expression measured on the two adjacent tissue sections as 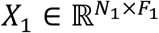 and 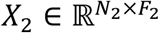, respectively, where the molecular sets are non-overlapping across the two sections. Here, the constant *N*_1_ and *N*_2_ denote the numbers of cells captured within the respective sections, whereas the constant *F*_1_ and *F*_2_ denote the numbers of profiled molecular features in *X*_1_ and *X*_2_, respectively. Throughout the following methodological formulations, we utilized *X* as a unified notation when no explicit distinction was required of the two sections, indicating that the corresponding computational operation was applied to both sections. This identical convention was consistently extended to other algorithmic variables and parameters unless stated otherwise.

The architecture of SpaWeaver (Fig. 1c) comprises four major components: (1) an H&E foundation model, (2) a graph Transformer module, (3) a latent feature aligner, and (4) dual omics predictors. Within each individual tissue slice, SpaWeaver first deploys an H&E foundation model to extract morphologically informative features from either real or virtual histology images. Then, a graph Transformer module integrates multi-scale spatial context across the spatial cell graph to generate spatially aware cellular morphological embeddings. To accommodate weakly anchored data, the latent feature aligner is specifically designed to mitigate domain shifts in the learned representations. Finally, each omics-specific predictor infers the corresponding omics profiles from the encoded cellular embeddings. Together, these components enable SpaWeaver to establish a direct, end-to-end computational bridge between tissue morphologies and molecular profiles across different omics. Detailed mathematical and algorithmic formulations of each constitutive module are provided in the subsequent sections.

### H&E foundation model

To encode cellular morphological information into a latent representation space, SpaWeaver first extracts square image patches of size *r* × *r* centered on the coordinates of each cell. The resulting collection of these histology patches is denoted as *P* ∈ ℝ^*N*×*r*×*r*×3^. These patches were subsequently fed into a pretrained pathology foundation model to obtain histological feature embeddings. In this study, we deployed UNI^19^ as the default pathology encoder owing to its visual representation capability across diverse computational pathology tasks. After feature extraction, the histology patch of each cell is transformed into a high-dimensional vector *H* ∈ ℝ^*N*×*M*^, which served as the input for subsequent computational modules. Here, the constant *M* denoted the embedding dimensionality of the H&E foundation model, where *M* = 1024 for the UNI architecture. Accordingly, the complete feature matrices derived from the two adjacent tissue sections are denoted as *H*_1_ and *H*_2_, respectively.

### Graph Transformer module

To capture complex spatial relationships among heterogeneous cells, we constructed a spatial proximity graph 𝒢 = (𝒱, ℰ) for each tissue section based on cellular coordinates, where the set 𝒱 = {*v*_1_, *v*_2_, ⋯, *v*_*N*_} denotes the cells (nodes) and the set ℰ represents the intercellular spatial connections (edges). For each cell, undirected edges were established between itself and its nearest spatial neighbors. Let *A* ∈ ℝ^*N*×*N*^ denote the adjacency matrix of graph 𝒢. The initial cellular feature matrix input to this model was denoted as *H* ∈ ℝ^*N*×*M*^, where each feature vector corresponds to the histological embedding of a cell extracted by the H&E foundation model. After constructing the spatial proximity graph, we employed a graph Transformer architecture to model multi-scale microenvironmental dependency for each cell. Unlike conventional graph neural networks that rely on localized message passing, this graph Transformer leverages a global attention mechanism, enabling each cell to aggregate information from long-range spatial contexts while preserving local spatial context within a single forward pass. Inspired by NAGphormer^20^, we adopted a modified graph Transformer-based architecture whose core components comprise a neighborhood tokenization strategy and a neighborhood message-passing mechanism. The former represented each individual cell as a discrete token sequence containing multiple neighborhood contexts, whereas the latter leveraged the Transformer layers to model multi-scale spatial dependencies.

Specifically, we first employed a simplified graph convolution network^66^ to generate cellular embeddings that integrated *ϰ*-hop neighborhood information, and *ϰ* denotes the predefined maximum hop distance. This process can be formalized as follows:

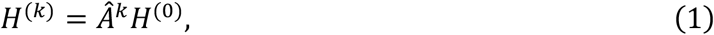

where the index *k* ∈ {0, 1, …, *ϰ*}, *H*^(0)^ = *H* denotes the initial morphological feature matrix, and *H*^(^*ϰ*^)^ signifies the spatially aggregated cellular embedding matrix at the *ϰ*-th hop. The operator 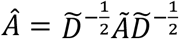 denotes the symmetrically normalized adjacency matrix, obtained by adding self-loops to the spatial proximity graph (*Ã* = *A* + *I*) and rescaling each edge weight by the inverse square root of the degrees 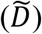 of the two connected nodes.

After the propagation process formulated in Eq. (1), the resulting embeddings are subsequently organized into a structured sequence encoding cellular microenvironmental context, denoted as 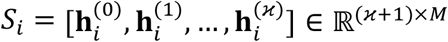. Accordingly, the sequences for all cells across the entire tissue section can be compactly expressed as *S* = [*S*_1_, *S*_2_, …, *S*_*N*_] ∈ ℝ ^*N*×(ϰ+ 1)× *M*^. Notably, the propagation operation in Eq. (1) is entirely parameter-free, which allows these multi-hop neighborhood embeddings to be precomputed prior to model training, thereby substantially improving computational scalability and mitigating memory bottlenecks during optimization.

Subsequently, a projection function *ϕ*(⋅) was applied to reduce the dimensionality of the cell sequence embeddings *S* before feeding into Transformer encoder:

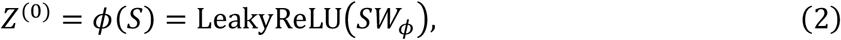

Where the matrix *Z*^(0)^ ∈ ℝ^*N*×(ϰ+ 1)× *d*^ denotes the projected latent representations, *W*_*ϕ*_ ∈ ℝ^*M*×*d*^ represents the trainable weight matrix utilized for dimension reduction, and LeakyReLU(⋅) signifies the leaky rectified linear unit activation function^67^ that introduces non-linearity into the projection function.

We fed *Z*^(0)^ into a Transformer encoder consisting of stacked Transformer layers, each comprising a multi-head self-attention (MSA) module^28^ and a position-wise feed-forward network (FFN). For clarity of exposition, we describe the self-attention mechanism using a single-head formulation. Specifically, *Z*^(0)^ is first projected into query, key, and value subspaces, denoted as *Q, K, V*, respectively. The queries and keys are then utilized to compute the scaled dot-product attention, followed by a row-wise softmax(⋅)^68^ normalization to obtain the attention matrix. This matrix is subsequently multiplied by the values to generate the updated hidden embeddings *Z*′ ∈ ℝ^*N*×(ϰ +1)× *d*^:

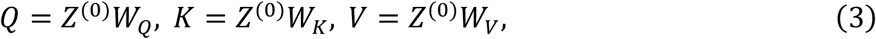

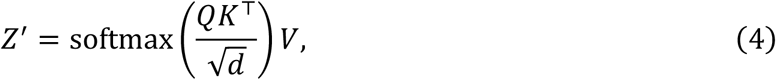

where *W*_*Q*_, *W*_*K*_, *W*_*V*_ ∈ ℝ^*d* × *d*^ denote the trainable projection weight matrices across the respective subspaces, and the scalar *d* represents the dimensionality of the query and key vectors.

MSA extends the single-head attention mechanism by projecting the query, key, and value matrices described in Eq. (3) into multiple independent subspaces, denoted as *Q*^(𝒽)^, *K*^(𝒽)^, and *V*^(𝒽)^, respectively. The MSA operation is defined as:

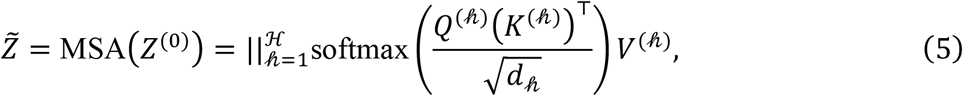

where *d*_𝒽_ = *d*/ℋ denotes the feature dimensionality of each attention head, ℋ represents the total number of attention heads, and “||” denotes the concatenation operation.

MSA enables the model to concurrently aggregate complementary relational information across multiple representation subspaces. The resulting head-wise embeddings matrices were concatenated along the feature dimension to seamlessly reconstruct the final latent representation matrix 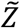.

Following the MSA module, the resulting representation matrix 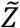 is processed using layer normalization^69^, and feed into a position-wise FFN module. This FFN component was composed of two consecutive linear transformations partitioned by a non-linear GELU(⋅) activation function^70^, which was formalized as follows:

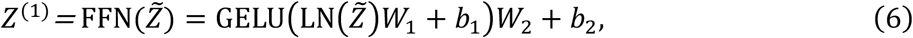

where the operator LN(⋅) denotes layer normalization, the matrices *W*_1_, *W*_2_ ∈ ℝ^*d*×*d*^ denote trainable weight matrices, and *b*_1_, *b*_2_ are bias terms.

The above equations describe the computation within a single Transformer layer. When stacking multiple Transformer layers to construct a deep architecture, the overall computation stream was expressed as follows:

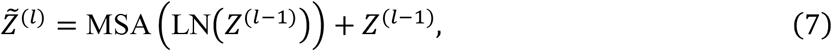

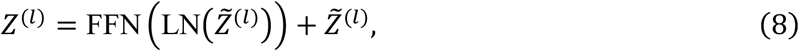

where the *l* ∈ {1, ⋯, *L*} indexes the Transformer layers, and *L* represents the total number of stacked layers.

After passing through *L* graph Transformer layers, we obtained the latent representation *Z*^(*L*)^ ∈ ℝ^*N*×(ϰ +1)× *d*^, which encodes both the intrinsic features of the target cells and the structural context of their surrounding multi-scale spatial microenvironments. Given that neighboring cells at different topological hops may exert distinct influences on the target cell depending on their spatial proximity, we further employed an attention-based readout function adaptively aggregating the token sequence and derived the final compact cellular embedding matrix *Z* ∈ ℝ^*N*×*d*^. This readout process can be formulated as follows:

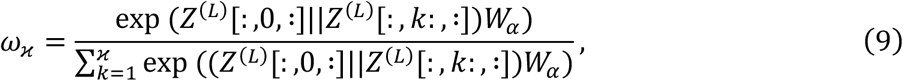

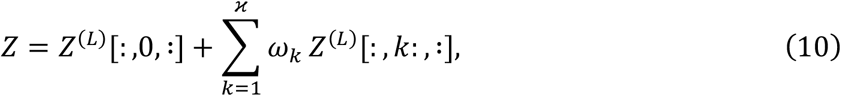

where *ω*_*k*_ denotes the attention coefficient and *W*_*α*_ represents the trainable weight matrix utilized for subspace transformation during aggregation.

For simplicity, the comprehensive morphological representations of the two distinct tissue slices, that is, the initial feature matrices *H*_1_ and *H*_2_, after passing through the graph Transformer encoder and the subsequent readout module, were formally denoted as *Z*_1_ and *Z*_2_, respectively.

### Latent feature aligner

Because the two weakly anchored adjacent H&E-stained images were acquired under different experimental protocols, or because the virtual H&E image differs from real H&E image, the extracted latent embeddings predominantly encoded localized cellular morphology heavily corrupted by these non-biological variations, thereby leading to a noticeable domain shift between *Z*_1_ and *Z*_2_.

To effectively mitigate the domain shift in the shared latent space, we employed maximum mean discrepancy (MMD)^29^ as a non-parametric distribution alignment metric. MMD maps samples into a high-dimensional reproducing kernel Hilbert space (RKHS) via specific kernel functions, thereby enabling the statistical discrepancies between the two slices to be quantitatively measured.

In our setting, given two cell embedding sample sets 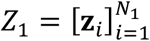 and 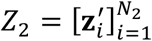 derived from the adjacent tissue slices, drawn from distributions ℙ and ℚ, respectively, the theoretical MMD between the two distributions is defined as follows:

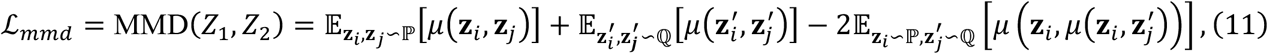

where *μ*(⋅,⋅) denotes a kernel function (e.g., a Gaussian kernel 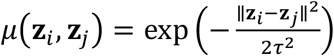, where *τ* is the bandwidth parameter), and 𝔼[⋅] represents the mathematical expectation operator. In practice, these continuous expectations are approximated using empirical sample means over the available cell populations, yielding the following simplified discrete formulation:

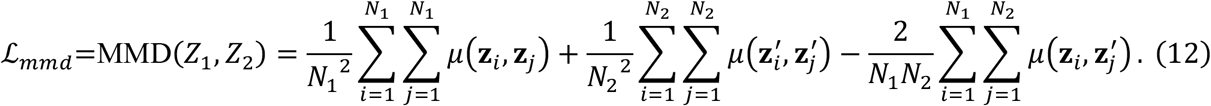

By minimizing the MMD loss defined in Eq. (12), we aligned the two domain-shifted cellular morphological representations within the shared latent space, facilitating the subsequent omics prediction module.

In the cross-resolution setting, ℒ_*mmd*_ alignment is conducted at the lower-resolution level. Taking the Visium–CODEX data integration as representative paradigm, SpaWeaver generated pseudo-spots on the high-resolution CODEX tissue section, with a specified diameter of 55 *μ*m to physically match the spot size of the Visium array. To guarantee that all CODEX cells were comprehensively subjected to the distribution constraint enforced by the MMD loss, neighboring pseudo-spots were allowed to overlap with each other, rather than adhering to the conventional 100 *μ*m center-to-center spacing of the standard Visium layout. This procedure yielded a pseudo-spot-by-cell assignment mapping matrix, which was subsequently utilized to aggregate the single-cell-level CODEX representations into pseudo-spot-level representations prior to computing MMD loss against the Visium spot-level representations. Crucially, this assignment mapping matrix was not deployed in the subsequent decoder-based mean square error loss. Instead, the final multi-omics decoder was trained rigorously on the original cell-level outputs. This decoupled optimization strategy explicitly encouraged the model to faithfully retain highly fine-grained spatial omics variations, preventing the network from learning only spatially smoothed, pseudo-spot-level blurred patterns.

### Omics predictor

After obtaining domain-aligned cellular morphological representations *Z*_1_ and *Z*_2_, SpaWeaver can optionally incorporate additional metadata to further enrich the representations used for omics prediction. Taking cell-type annotations as an example, SpaWeaver encodes each cell type as a learnable embedding vector, forming a cell-type embedding dictionary. For cell-resolution patches, the annotation vector is directly given by the embedding corresponding to the annotated cell type. For spot-resolution patches, the annotation vector is computed as a weighted average of the cell-type embedding dictionary, where the weights are given by the deconvolved cell-type proportions. The resulting annotation vector is concatenated with *Z*_1_ and *Z*_2_ before being passed to the omics decoders. Because this metadata-enhancement step is optional, we continue using *Z*_1_ and *Z*_2_ in the following sections, unless otherwise specified.

To establish a high-fidelity mapping between the cellular morphological features and their corresponding molecular profiles, we employed two independent omics predictors, denoted as *χ*(⋅) and *ξ*(⋅), respectively. These predictors were designed to map the aligned cellular representations *Z*_1_ and *Z*_2_ back into their respective molecular profile spaces, which can be formalized as follows:

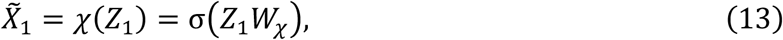

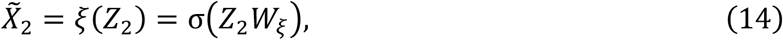

where 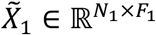 and 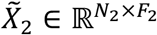 denote the reconstructed omics profile matrices, and *W*_*χ*_ and *W*_*ξ*_ are trainable weight matrices.

To enforce strict supervision during the joint training phase, the reconstruction objective minimized the dataset-specific mean squared error calculated between the predicted and measured omics profiles, which guided the model via the following formulation:

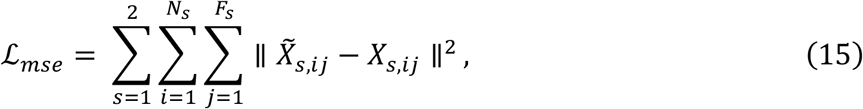

where *s* is the slice index, *i* is the index for cells and *j* is the index for molecular entities, e.g., genes and proteins.

### Model optimization

To simultaneously achieve high-fidelity multi-omics reconstruction and robust cross-slice domain shift removal, the reconstruction and MMD alignment losses were jointly optimized using a multi-task learning framework. The final comprehensive objective function was formulated as follows:

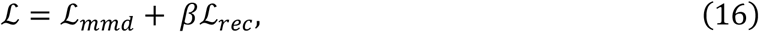

where ℒ denotes the total joint loss, and *β* represents a hyperparameter weighting coefficient that dynamically controls the trade-off contribution of the MMD distribution alignment term during the global backpropagation process.

### Implementation details

During spatial proximity graph construction, we connected each cell to its 7 spatially nearest neighbors based on Euclidean distance. The graph Transformer encoder adopted a shallow single-layer, two-head architecture (*L* = 1, ℋ = 2), with the latent dimensionality fixed to *d* =128. For the latent feature aligner, the bandwidth parameter of the Gaussian kernel utilized in the MMD computation was fixed to *τ* = 1. The global loss weighting coefficient that balances the multi-task objectives was set to *β* = 0.01. These parameters were fixed across the datasets used in this study. Ablation studies regarding core model components and extensive parameter sensitivity analyses for the choice of cellular H&E image patch radius, number of neighboring cells, and number of hops were systematically presented in Supplementary Fig. 7.

The entire SpaWeaver was trained end-to-end using the Adam^71^ optimizer with a learning rate of 0.001 for 500 epochs. All experiments were conducted using PyTorch^72^ (version 2.3.1) and Python (version 3.8) environments deployed on multiple NVIDIA L40 GPUs with 48 GB memory.

## Data

### Spatial transcriptome data

We applied platform-specific quality control procedures for different ST technologies, followed by a unified preprocessing pipeline. All these steps were implemented using the Scanpy^73^ package.

#### Quality control

For Xenium datasets, cells with fewer than 50 total gene expression counts were removed. For Visium HD datasets, analyses were performed at 8 *μ*m resolution. Bins with fewer than 50 total gene expression counts were filtered out, and bins with more than 20% mitochondrial gene expression were excluded.

#### Preprocessing

Raw gene expression counts were normalized using *scanpy*.*pp*.*normalize_total* followed by log-transformation with *scanpy*.*pp*.*log1p*. For Visium, Visium HD, Xenium 5K datasets, the top 500 spatially variable genes were selected using Moran’s *I* statistics. For Xenium datasets, no gene filtering was applied, as the targeted gene panel comprises a limited and predefined set of genes.

#### Simulation of stain variability

To simulate staining variability across experiments, we applied perturbation on H&E images. Specifically, the hematoxylin and eosin channels were independently perturbed using randomly sampled multiplicative scaling factors from [1 − *α*, 1 + *α*] and additive offsets from [−*β, β*], with *α* = *β* = 0.2. The D (background) channel was not perturbed. This perturbation introduces moderate staining variation while preserving underlying tissue morphology.

### Spatial proteomics data

#### Preprocessing

For spatial proteomics data, protein intensities were normalized using a per-cell geometric mean scaling and subsequently log-transformed, as commonly used in prior studies^74^.

#### Virtual H&E generation

To overcome the lack of H&E images, we generated virtual RGB images using the TissueTag package^14^, which implements a method adapted from Simonson *et al*.^75^ to construct virtual H&E images from two user-defined fluorescent channels.

#### Annotation

For each cell, marker-based scores were calculated for each candidate cell type using curated marker proteins. Cells were then ranked according to their scores for each cell type, and labels were assigned by selecting the highest-scoring cells until the predefined proportion for that cell type was reached. These proportions were estimated from the cell-type annotations of the adjacent spatial transcriptomics section.

### H&E image denoising

We observed periodic stripe-like artifacts in cell-level H&E embeddings from several low-quality H&E images, including the COAD and OV samples in the SPATCH dataset, despite these artifacts being barely visible in the raw images. To reduce their effect on model training, we applied sinusoidal filtering to each embedding dimension along the spatial x and y axes. Barcode-based data used the original barcode coordinates, whereas cell-resolution data were spatially binned before frequency-domain artifact detection and sinusoidal component removal. For the OV dataset, an additional autoencoder-based denoising step was applied to remove residual nonlinear noise. The source code is publicly available in the SpaWeaver package.

### Evaluation

In this study, three evaluation metrics, including Pearson correlation coefficients (PCC), correlation matrix distance (CMD), and the structural similarity index measure (SSIM), were utilized for quantitatively assessing the model’s performance.

**PCC** measures the strength of the linear relationship between two variables, and is defined as follows:

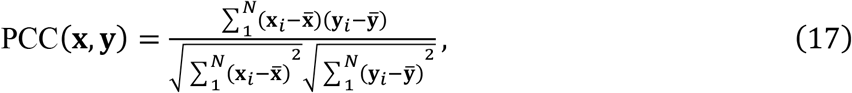

where **x, y** ∈ ℝ^*N*^ represents the expression of a specific gene across *N* cells, respectively. 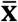 and 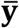 represent the mean values of **x** and **y**.

**SSIM** is adopted to assess the similarity between the ground-truth and predicted spatial structures of gene expression, where higher values indicate stronger structural agreement. In this work, SSIM is further employed to quantify the structural similarity of cell graphs constructed from spatial information. The formulation is given as follows:

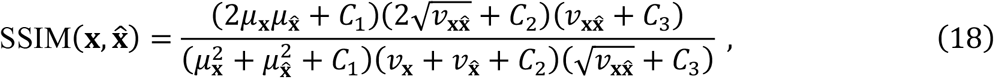

where, **x** and 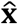 denote the measured and predicted gene expression, respectively. The terms *μ*_**x**_ and 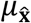 represent the spatially Gaussian-filtered means of **x** and 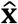, while *v*_**x**_ and 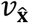 correspond to their spatially Gaussian-filtered variances. The term 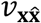 denotes the spatially Gaussian-filtered covariance between **x** and 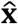. The constants *C*_1_, *C*_2_, and *C*_3_ are predefined to stabilize the computation, where *C*_1_ = (0.01(max(**x**) − min(**x**)))^2^, *C*_2_ = (0.03(max(**x**) − min(**x**)))^2^, and 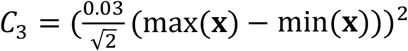.

**CMD** provides a general measure for quantifying the discrepancy between two correlation matrices, *R*_1_ and *R*_2_, with smaller values indicating closer agreement. The CMD is defined as follows:

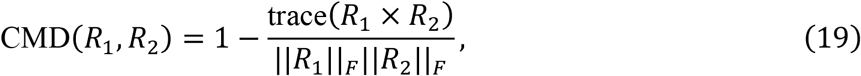

where trace(⋅) denotes the trace of a matrix, and ∥ ⋅ ∥_*F*_ represents the Frobenius norm of the matrix.

### Spatial domain identification

In this study, spatial domain identification is performed in two distinct contexts, both implemented using CellCharter^44^, but serving different purposes. In the quantitative evaluation, spatial domain identification is used to indirectly assess whether the predicted gene expression preserves the underlying biological structure. Specifically, we applied CellCharter independently to the observed gene expression and to the predicted gene expression. The spatial domains derived from the observed gene expression are treated as the ground truth, and the agreement between the two domain partitions is quantified using the ARI. In the qualitative analysis, we jointly input the gene expression profiles predicted by each method across the two tissue sections into CellCharter for integrated spatial domain identification. In addition, we attempted to mitigate domain-shift effects in affected methods using scVI-based batch correction^45^, directly relying on the scVI implementation provided in the CellCharter package.

### Usage of scIB package^34^

We evaluated batch effect removal performance using established metrics implemented in scIB, including batch correction metrics (iLISI, S-batch, kBET, ASW, and graph connectivity) and biological conservation metrics (cLISI, S-label, and kNMI). All metrics were computed on the concatenated gene expression profiles of the two predicted panels from both slices, followed by dimensionality reduction to the top 50 principal components. For computational efficiency, we down-sampled the data to a maximum of 100,000 cells before metric computation. The clustering step for kNMI was implemented using K-Means, whereas all other metrics were computed using the original implementations provided in the scIB package.

**scGraph**^35^ is used to evaluate whether the gene expression profiles predicted by a given algorithm preserve the underlying biological structure of the original data. Specifically, cell-type centroids are computed within each batch using trimmed means in PCA space, and pairwise distances between centroids are used to construct cell-type relational graphs. The consistency between graphs derived from predicted representations and a batch-wise consensus graph is then quantified using distance-weighted correlation metrics.

**GTE**^42^ is used to quantify batch effects in the predicted gene expression from a gene-centric perspective, assessing whether genes exhibit batch-specific biases across the two slices. A lower GTE value indicates better results. For a given gene *m*, the GTE value is defined as follows:

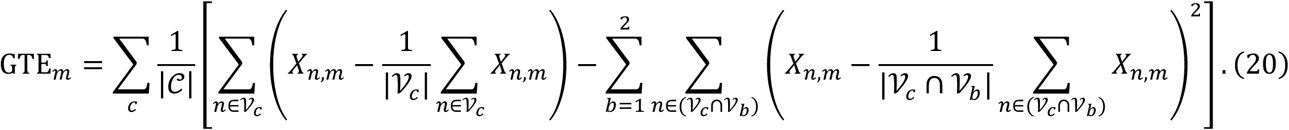

Here, *X* denotes the concatenated gene expression matrix obtained by stacking the predicted gene expression from the two slices, with 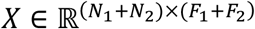. 𝒞 denotes the set of cell types, 𝒱 the set of all cells, 𝒱_*c*_ the subset of cells belonging to cell type *c*, and 𝒱_*b*_ the subset of cells belonging to batch *b*.

### Cell type defined spatial niche

For each cell, we first calculated the cell-type composition within a 50 *μ*m neighborhood. The resulting neighborhood cell-type composition vectors were then clustered using K-Means to define spatial niches.

### Cell–cell communication

In this study, cell–cell communication analysis was performed using CellPhoneDB^59^ as implemented in the LIANA package^76^. For each spatial niche, cell–cell communication was inferred separately using the *liana*.*method*.*cellphonedb()* function, with *resource_name = ‘consensus’*. Because protein expression measured by CODEX imaging represents continuous intensity values, we applied stringent filtering to the CellPhoneDB outputs. Specifically, only interactions satisfying all the following criteria were retained: cellphone_pvals < 0.05, ligand_pvals_adj < 0.05, and receptor_pvals_adj < 0.05.

### Resolution enhancement

We used CORAL^22^ to perform resolution enhancement on the pseudo-spots gene expression profiles generated by SpaWeaver after cross-resolution diagonal integration. Because SpaWeaver outputs overlapping pseudo-spots, we adapted the CORAL input format accordingly, and the modified code will be made publicly available on GitHub (see Methods “Code availability”). Spatial domains after resolution enhancement were identified by applying K-Means clustering directly to the representations learned by CORAL.

## Data availability

The datasets used in this study are publicly accessible. The Xenium Human Breast Cancer tissue dataset is available at https://www.10xgenomics.com/products/xenium-in-situ/preview-dataset-human-breast. The VisiumHD and Xenium sections in Human Colon Cancer P2 dataset is available at https://www.10xgenomics.com/platforms/visium/product-family/dataset-human-crc. The Xenium 5K and CODEX SPATCH human colon adenocarcinoma and ovarian cancer dataset is available at https://spatch.pku-genomics.org/.

## Code availability

Our code has been deposited in the MTS submission system and will be publicly available at https://github.com/KEAML-JLU/SpaWeaver. Detailed SpaWeaver tutorials will be released upon acceptance and have also been provided as Supplementary Note 1 in the MTS submission system. All processed datasets and analysis outputs generated in this study are available at https://zenodo.org/records/********.

## Author contributions

Z.Y., X.F., and R.G. conceived and supervised the study, secured funding, and provided analytical guidance. Y.L. and C.W. implemented the algorithm with assistance from Z.W. C.W. performed the main analyses with support from Y.L., Z.W. P.S. and Z.L. Y.L., J.L., X.W. and K.C implemented the benchmarking algorithms. C.W., J.L., and X.W generated the figures. K.C. built the code package and website. Z.Y., X.F., R.G., C.W., Y.L., and Z.W. wrote the manuscript with the input from all authors. Q.Z., D.Z., Z.H., and Y.D. provided helpful comments and assisted with manuscript polishing. All authors reviewed and approved the final version of the manuscript.

## Acknowledgements

Our work is supported by National Key R&D Program of China (No. 2023YFF1204800 (Z.Y.)), National Nature Science Foundation of China (No. 32470706 (Z.Y.), No. 62303119 (Z.Y), and No.62372209 (X.F.)), Shanghai Science and Technology Development Funds (No. 23YF1403000 (Z.Y.)), Chenguang Program of Shanghai Education Development Foundation and Shanghai Municipal Education Commission (No. 22CGA02 (Z.Y.)), Shanghai Municipal Science and Technology Major Project (No. 2023SHZDZX02 (B.Q.)), and Fund of Fudan University and Cao’ejiang Basic Research (No. 24FCA10(Z.Y.)).

## Inclusion & Ethics

Not relevant.

## Figure legends

**Fig. S1: Raw H&E images of human breast cancer dataset.a**. Raw H&E images on the serial slices in the human breast cancer dataset. **b**. Domain shift analysis of cellular histological representations extracted from the raw H&E images. Left: UMAP visualization showing minimal separation between the two serial sections before staining perturbation. Right: bar plots of quantitative metrics used to evaluate cross-section domain shift. Using the staining-perturbed condition as a reference, batch-effect-removal metrics indicated that the raw H&E-derived representations from the two sections were well mixed. Biological-conservation metrics further showed that the staining perturbation introduced minimal disruption to biologically relevant information encoded in the H&E images.

**Fig. S2: Benchmarking the impact of domain shift induced by weakly anchored data on all comparison methods. a**. UMAP visualization of the predicted gene expression by each method, colored by slice index. **b**. Top: Bat plots of mean GTE values across 313 genes with or without H&E style variation. Error bars represent the standard error of the mean (SEM). Bottom: Scatter plots of GTE values with or without H&E style variation colored by scatter density, with each point representing one of 313 genes.

**Fig. S3: Benchmarking spatial diagonal integration across nine methods on Xenium human breast cancer dataset with simulated H&E style variation using PCC. a**. Boxplot of PCC values computed between each methods predicted gene expression and measured expression with or without H&E style variation. Box plots show median, mean, quartiles and 1.5× interquartile range. **b**. Scatter plots of PCC values with or without H&E style variation colored by scatter density, with each point representing one of 313 genes.

**Fig. S4: Benchmarking spatial diagonal integration across nine methods on Xenium human breast cancer dataset with simulated H&E style variation using SSIM and CMD. a**. Scatter plots of SSIM values with or without H&E style variation colored by scatter density, with each point representing one of 313 genes. **b**. Bat plots of CMD values with or without H&E style variation. The two data points represent panel B imputed on slice 1 and panel A imputed on slice 2, respectively. Error bars represent the SEM.

**Fig. S5: Spatial visualization of gene expression predicted from each method on slice 2.** Measured expression is shown as reference.

**Fig. S6: Joint spatial domains identified on the two slices using the full panel (panel A + B) predicted by each method under the weakly anchored scenario.** Left: CellCharter jointly delineated spatial domain across the two slices, without applying scVI batch effect removal to the predicted expression. Right: CellCharter jointly delineated spatial domain across the two slices, with applying scVI batch effect removal to the predicted expression.

**Fig. S7: Ablation study and parameter sensitivity on Xenium human breast cancer dataset with simulated H&E style variation. a**. Bar plot of mean PCC values obtained by SpaWeaver and three ablated variants. **b**. Top: Bar plot of mean PCC values from SpaWeaver with different number of neighbors under 1 and 2 hops settings. Bottom: Bar plot of mean PCC values from SpaWeaver with different number of hops under 7, 15, and 50 neighbors settings. **c**. Bar plot of mean PCC values from SpaWeaver under different H&E patch radius. Error bars represent the SEM.

**Fig. S8: Annotation of COAD slices measured by Xenium and CODEX. a**. Cell type annotation on slice1 (Xenium). **b**. Cell type annotation on slice 2 (CODEX). **c**. Cell type annotation on slice1 (Xenium) with each cell type illustrated separately. **d**. Cell type annotation on slice2 (CODEX) with each cell type illustrated separately.

**Fig. S9: SpaWeaver predictions and downstream analysis on weakly anchored COAD data. a**. Left: Distribution of predicted protein abundance across cell types on slice 1, with measured protein expression patterns on slice 2 provided as reference. Right: Spatial visualization of predicted protein abundance on slice 1, with measured protein abundance on slice 2 shown as reference. **b**. Dot plot of *EPCAM* and differential gene between goblet cell and tumor epithelial. Spatial distribution of all epithelial cells, tumor epithelial and goblet cell are provided, with the spatial distribution of their marker genes expression.

**Fig. S10: Annotation of OV slices measured by Xenium and CODEX. a**. Left: Register spot distribution on H&E image. Right: Register cell distribution on virtual H&E image. **b**. Spatial distribution of cell type composition on slice 1, with each cell type shown separately. **c**. Left: Cell type annotation on slice 2. Right: Spatial distribution of cell type on slice 2, with each cell type shown separately.

**Fig. S11: SpaWeaver predictions and downstream analysis on weakly anchored OV data. a**. Super resolution gene expression generated by SpaWeaver workflow, with the measure expression on slice 1 shown as reference. The spatial abundance of corresponding coding protein is provided as reference if available. **b**. Spatial domain delineated using only measured proteomics on slice 2. **c**. Distribution of measured protein abundance across spatial domains on slice 2. **d**. Dot plot of marker genes for domain 1. **e**. Spatial distribution of marker genes of domain 1. **f**. Top: Selected ROIs enriched for domain 1 and other tumor-associated domains on slice 2. Bottom: Registration result of these ROIs on the H&E image from slice 1. **g**. Bar plot of the mean distance from cells in domain 2 to the nearest cell in other domains. Error bars represent the SEM.

**Supplementary Table 1: Benchmarking SpaWeaver and all comparison methods under panel diagonal integration with raw and perturbed H&E images using scIB metrics and scGraph.**

