## Supplementary Figure for "Robust integration of weakly anchored spatial multi-omics"

### Supplementary Figure 1

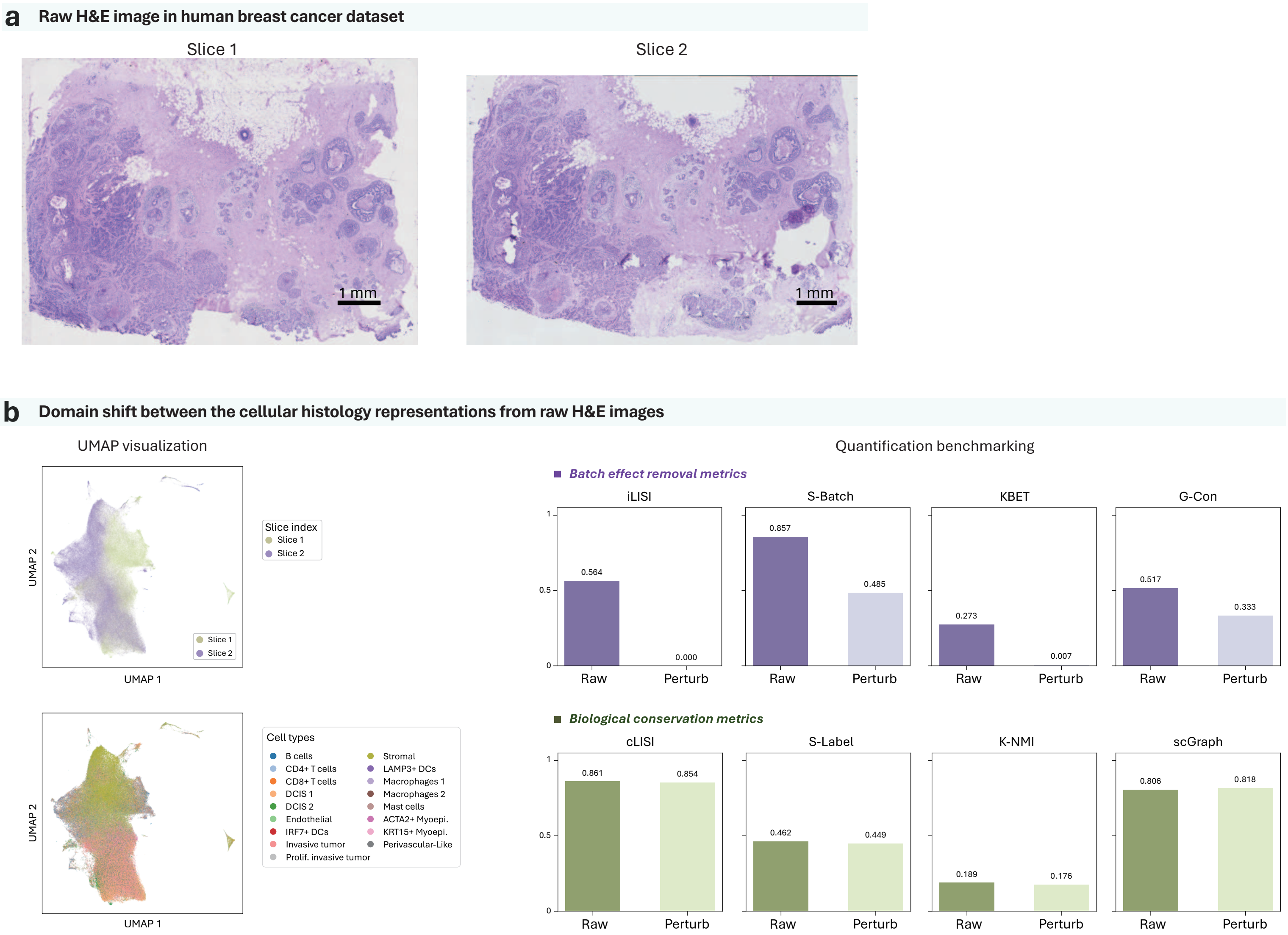

### Supplementary Figure 2

#### a UMAP visualization of the predicted gene expressions with perturbation from SpaWeaver and comparison methods

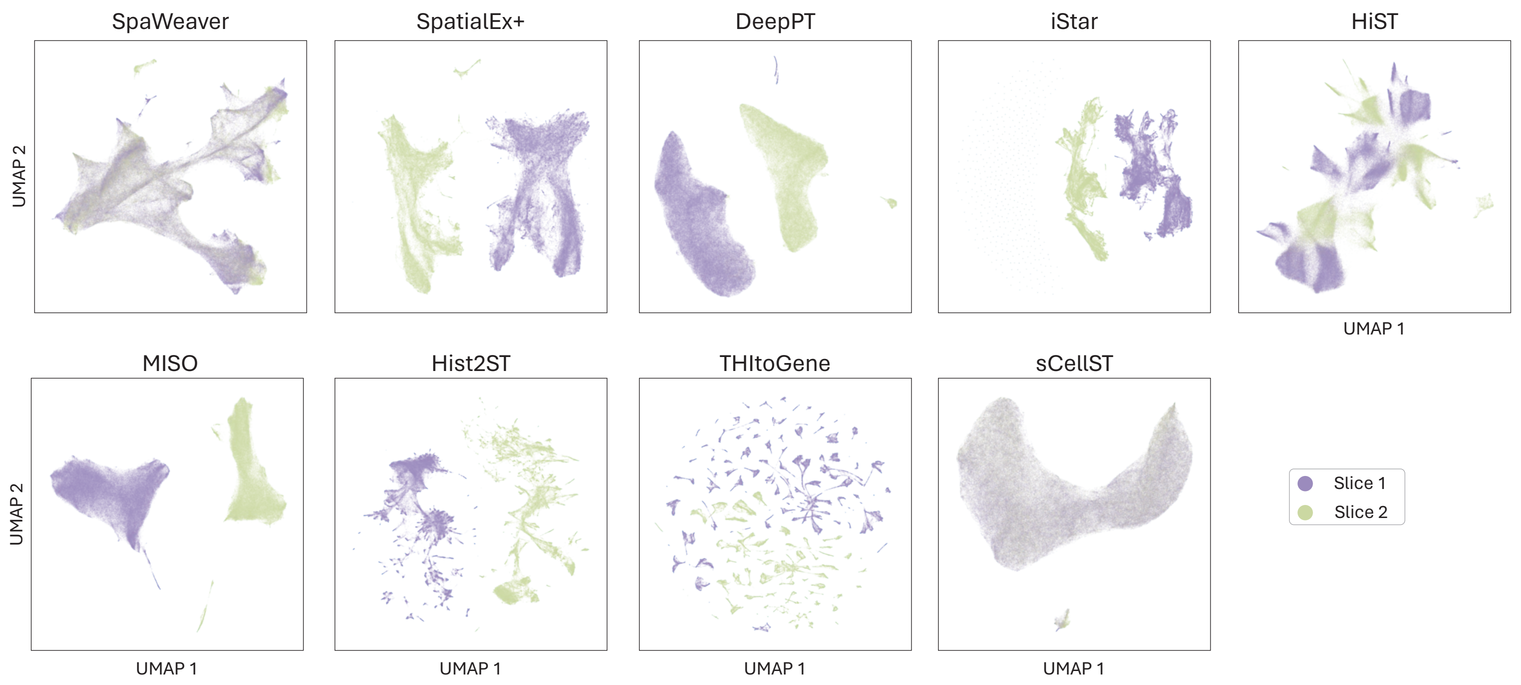


#### b GTE between measured and predicted gene expression by SpaWeaver and comparison methods

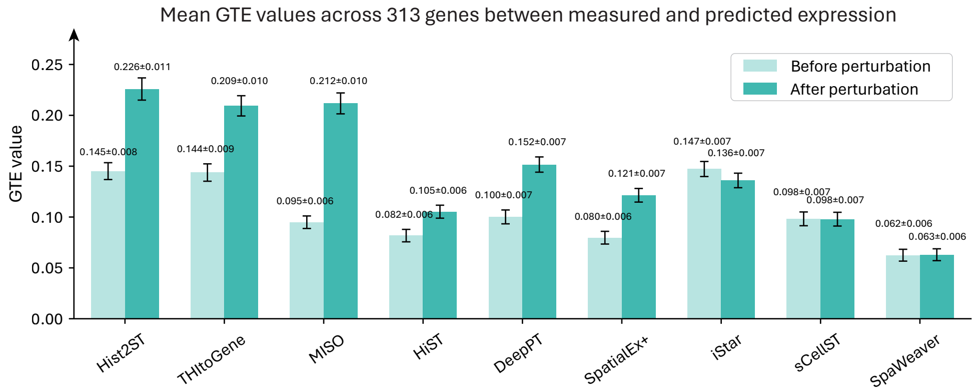


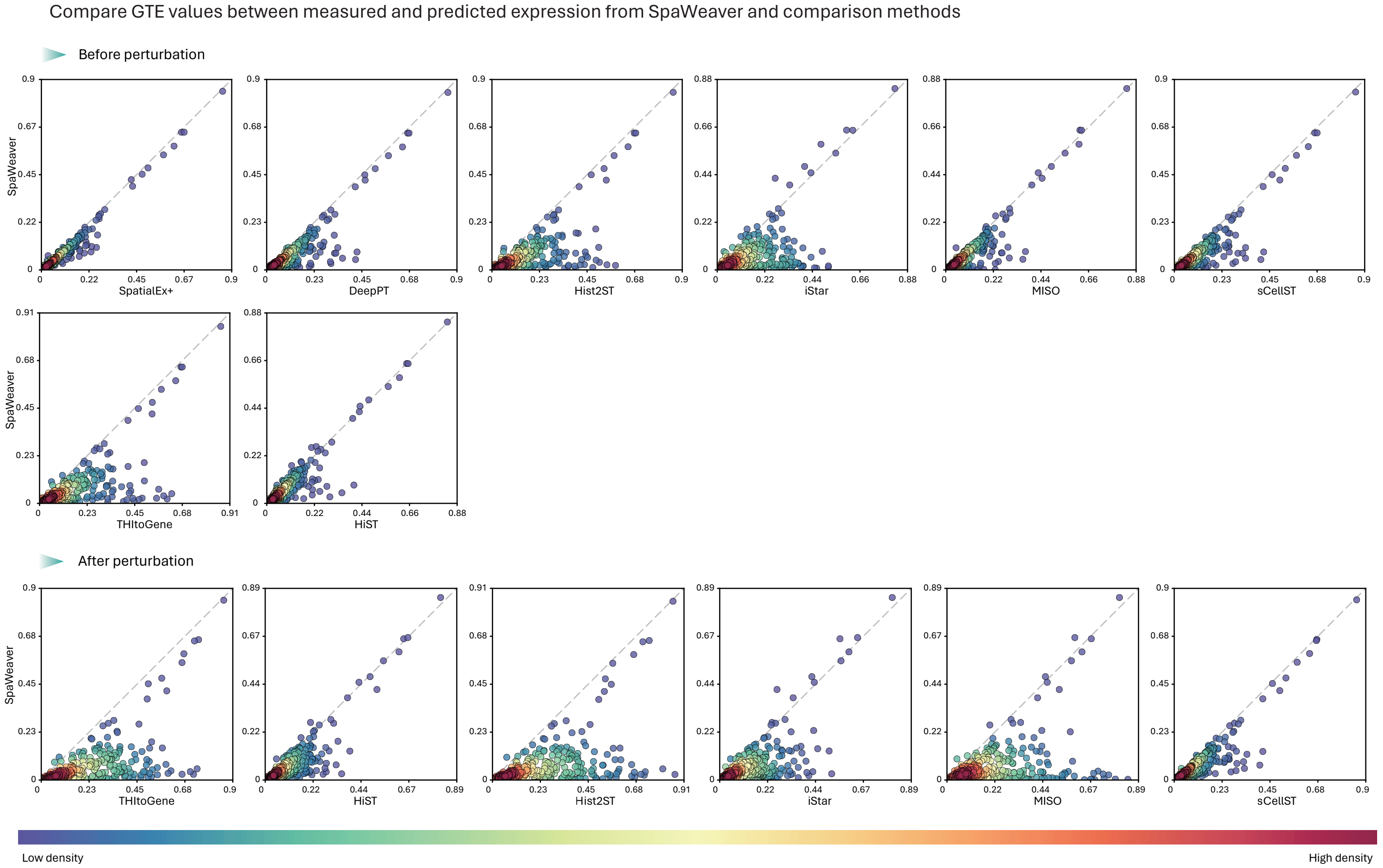

### Supplementary Figure 3

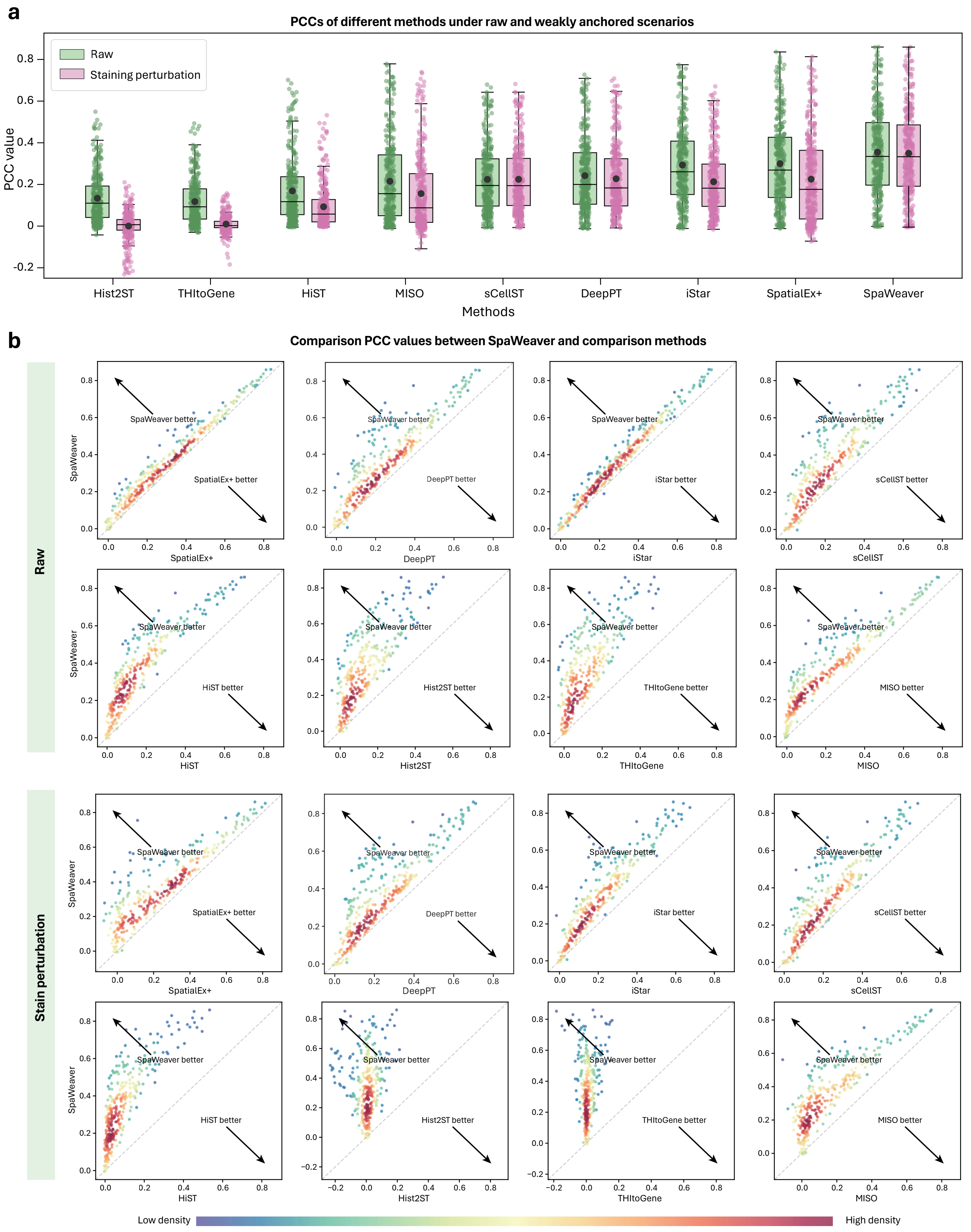

### Supplementary Figure 4

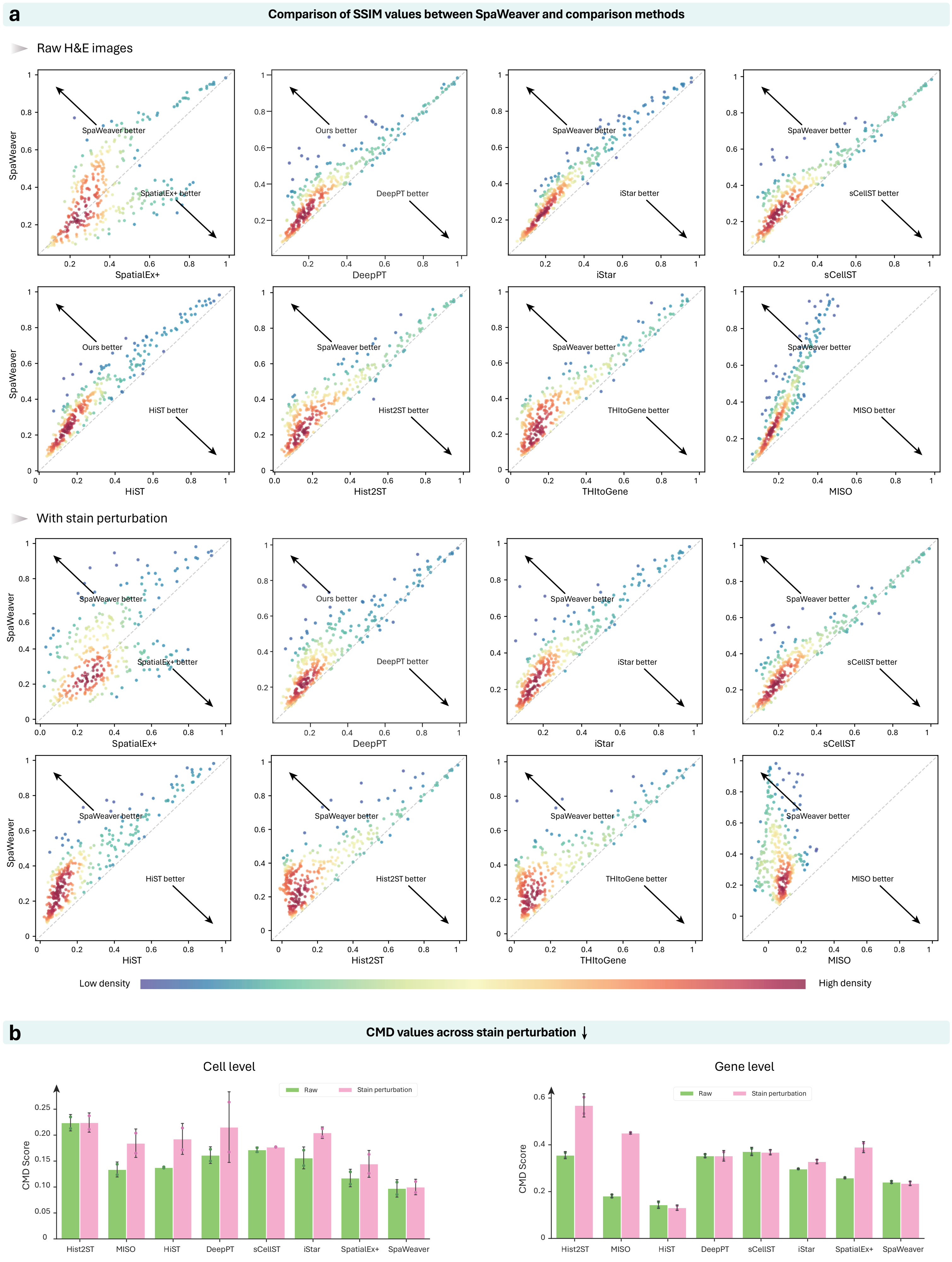

### Supplementary Figure 5

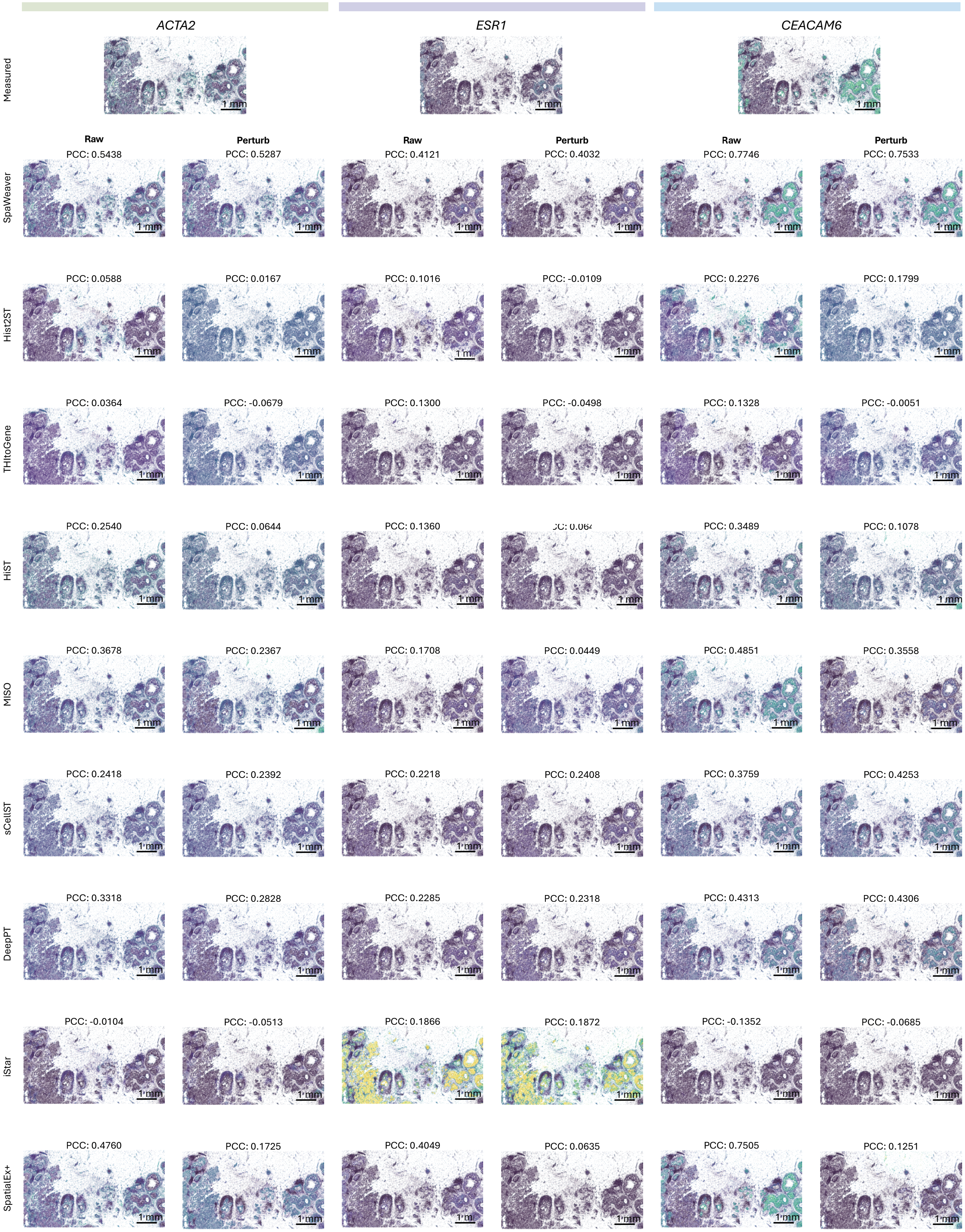

### Supplementary Figure 6

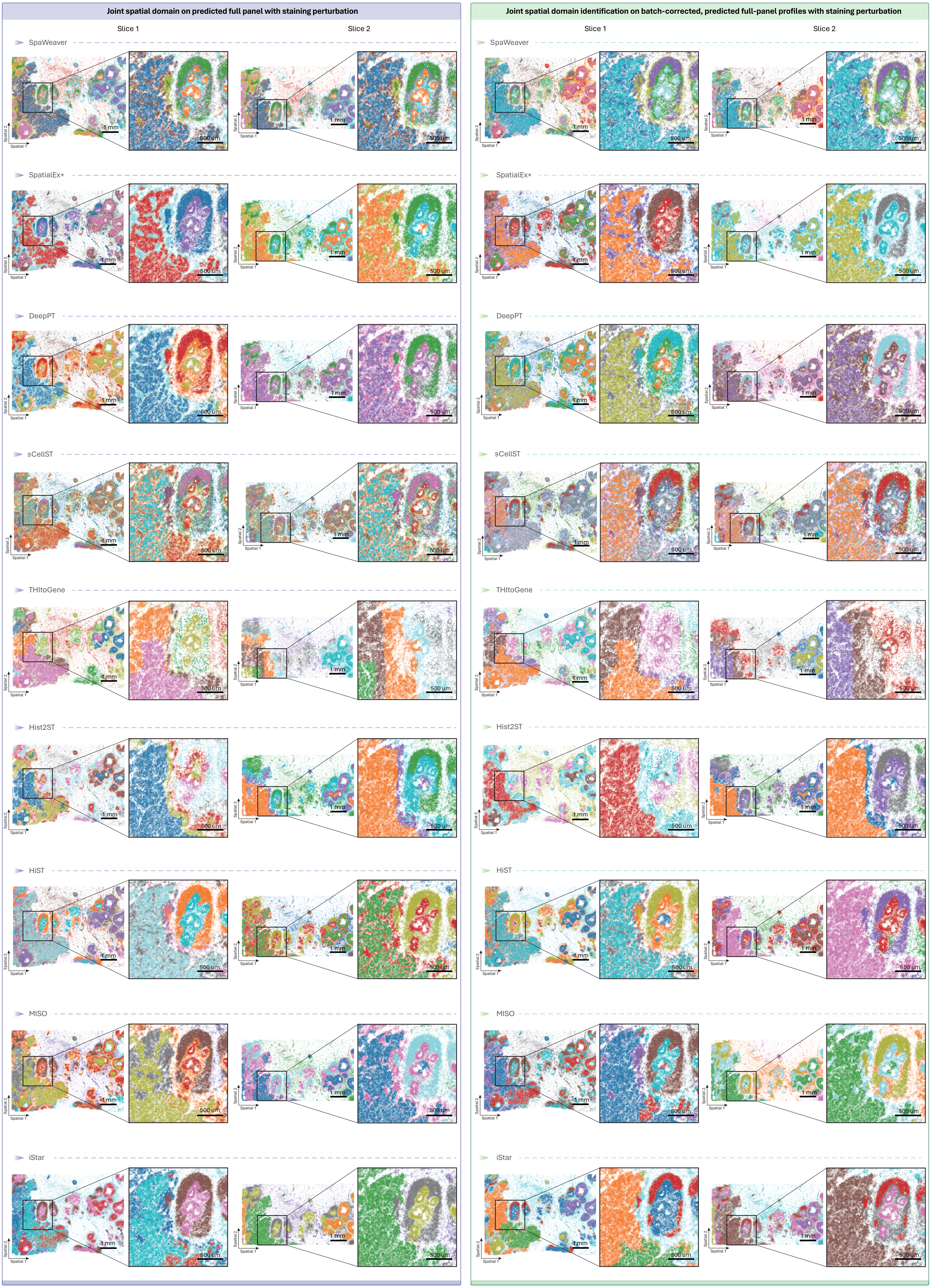

### Supplementary Figure 7

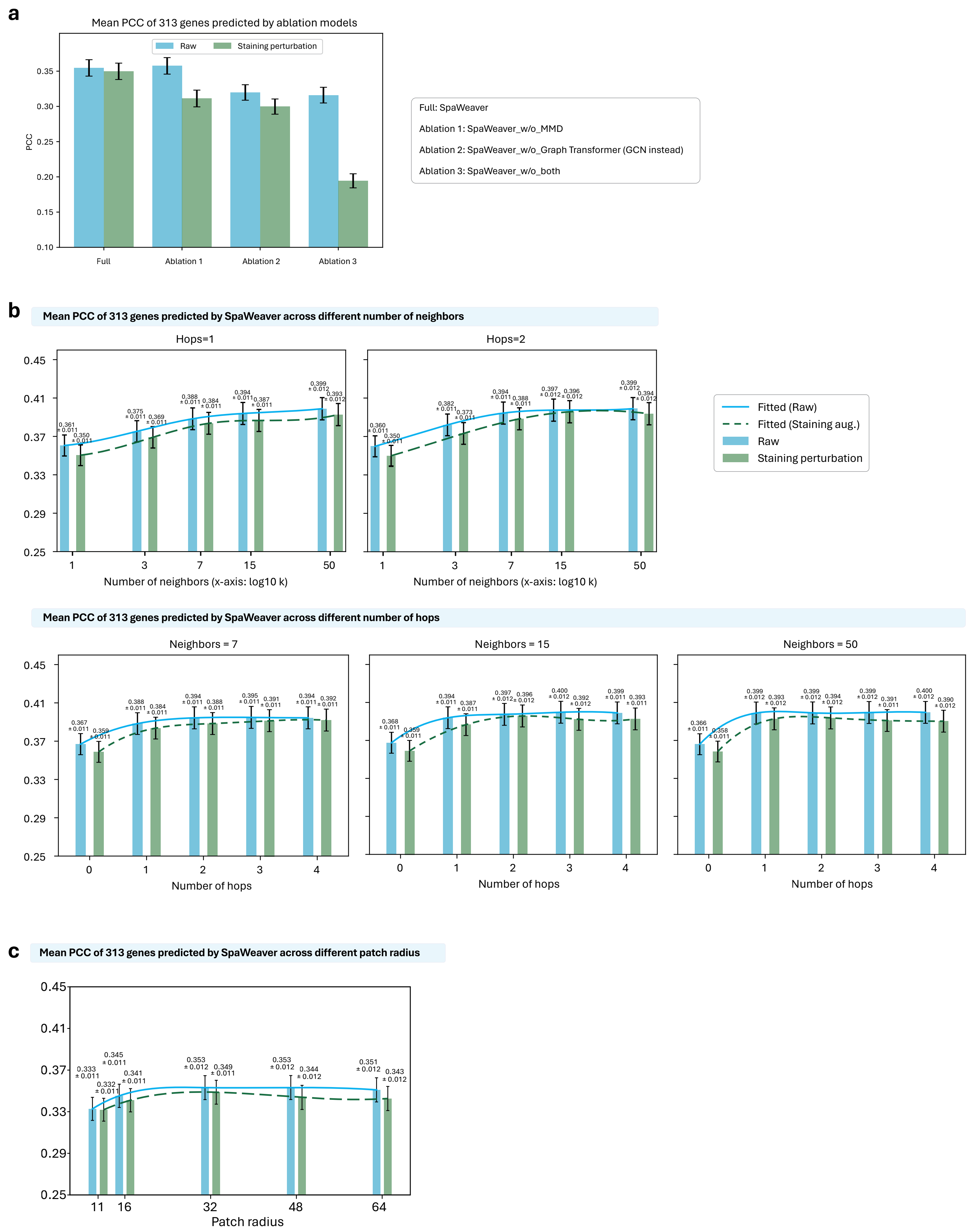

### Supplementary Figure 8

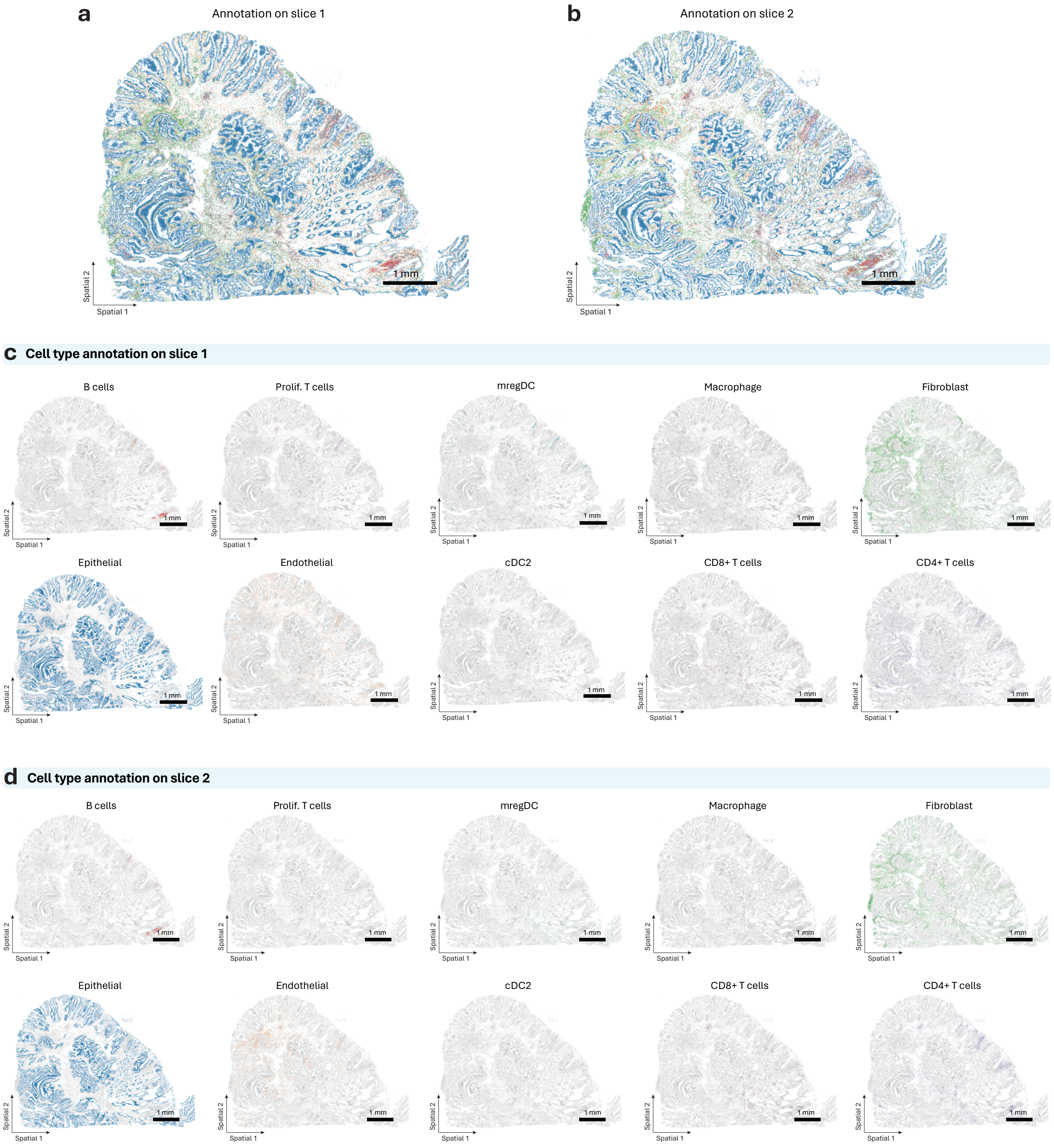

#### Supplementary Figure 9

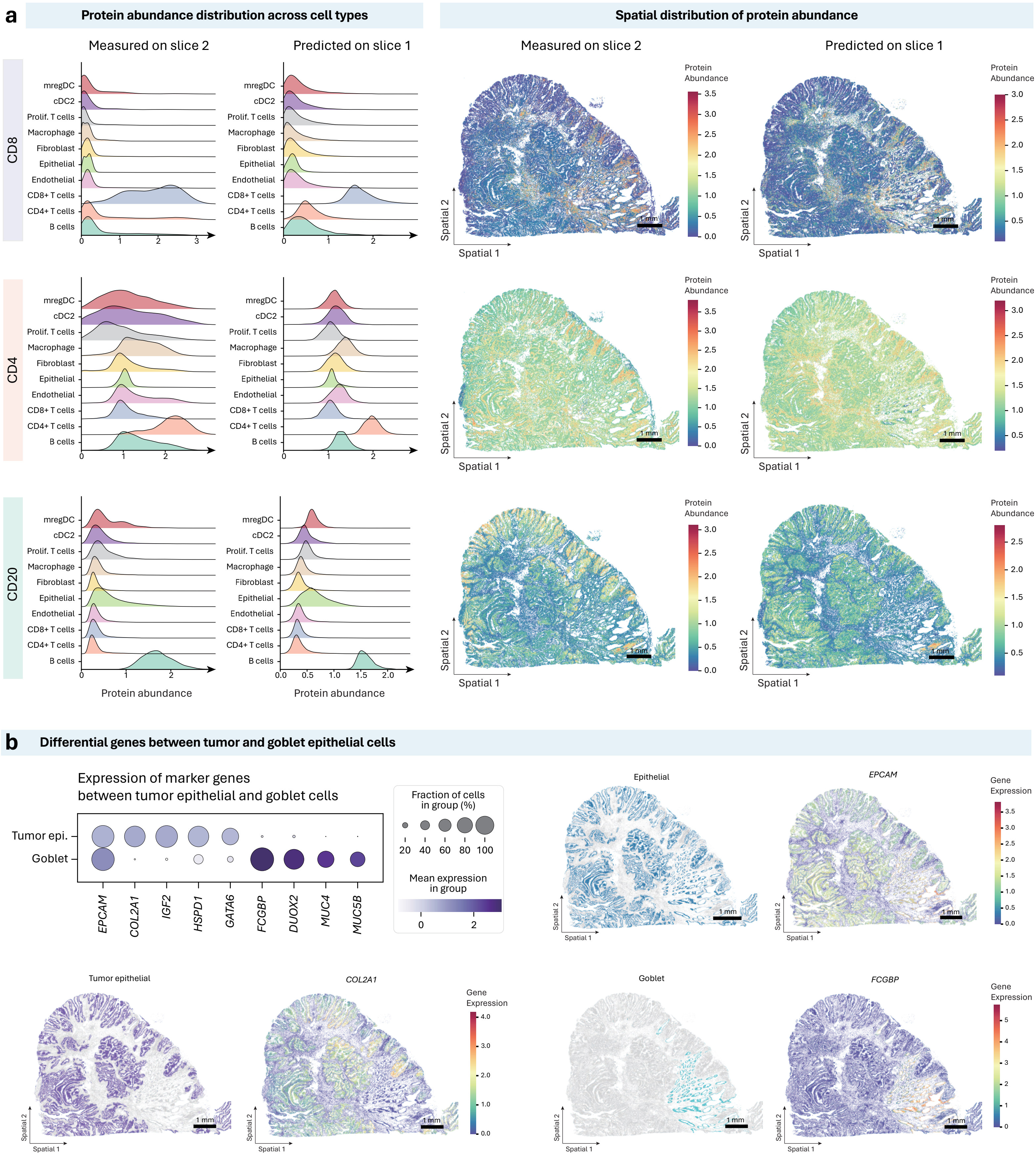

### Supplementary Figure 10

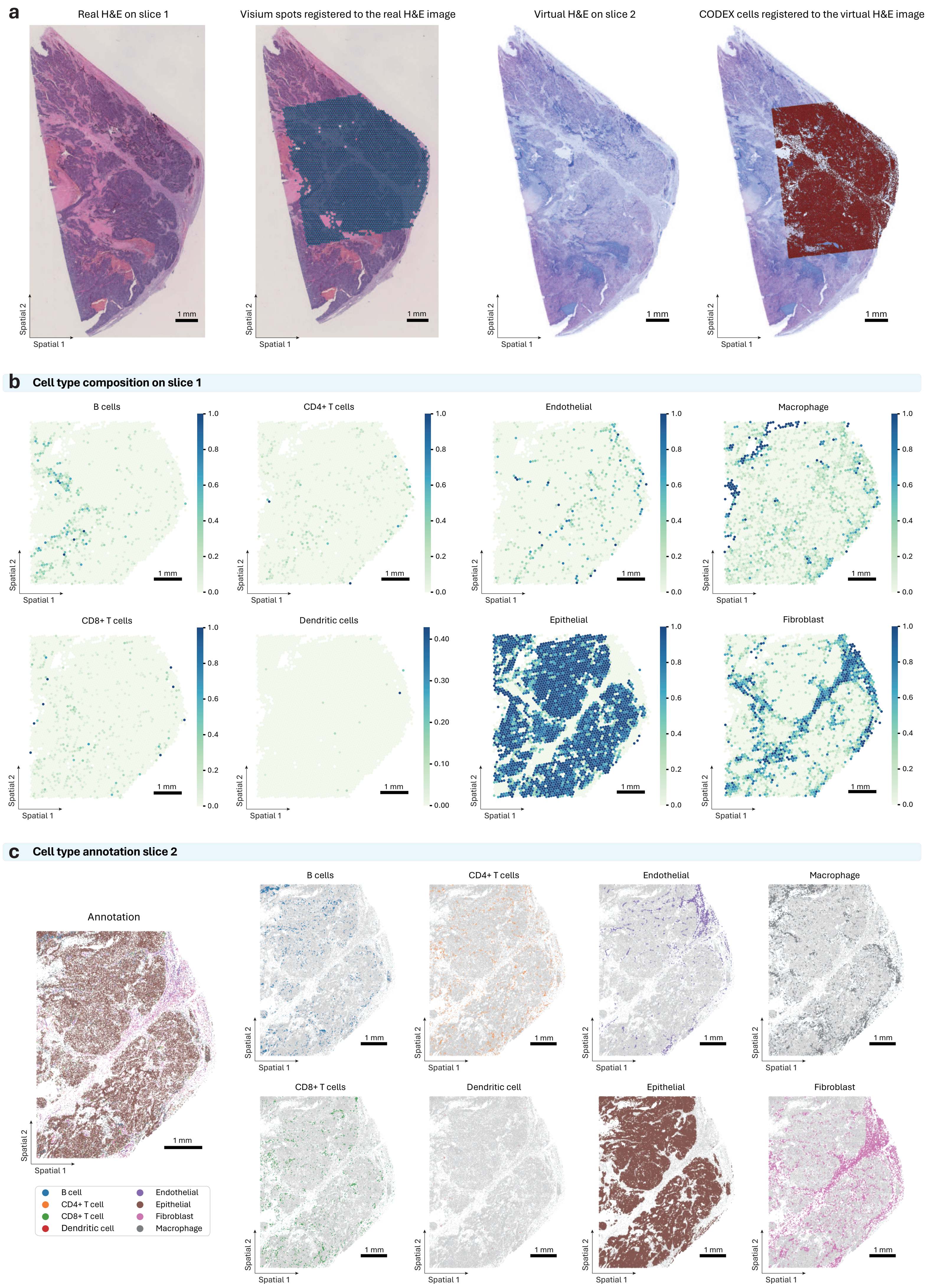

### Supplementary Figure 11

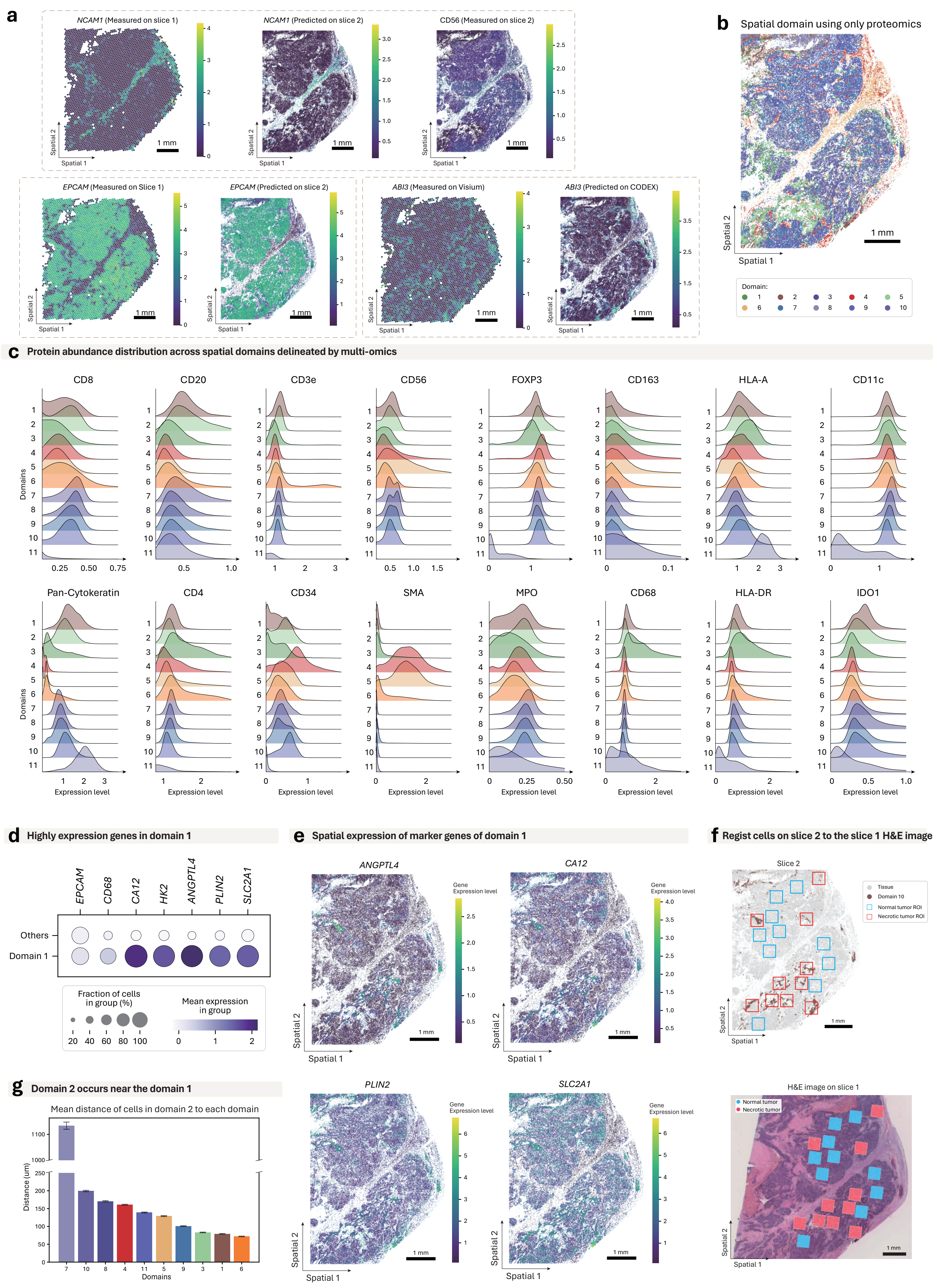

### Supplementary Table 1

**Raw H&E images**

| | iLISI | S-batch | KBET | G-Con | Batch overall | cLISI | S-label | K-NMI | scGraph | Bio. overall | Overall |
|---|---|---|---|---|---|---|---|---|---|---|---|
| SpaWeaver | 0.85 | 0.87 | 0.48 | 0.61 | 0.93 | 0.95 | 0.44 | 0.47 | 0.90 | 0.97 | 0.95 |
| DeepPT | 0.86 | 0.88 | 0.42 | 0.52 | 0.87 | 0.88 | 0.42 | 0.25 | 0.85 | 0.81 | 0.84 |
| SpatialEx+ | 0.68 | 0.85 | 0.29 | 0.54 | 0.75 | 0.93 | 0.41 | 0.40 | 0.85 | 0.90 | 0.83 |
| Hist2ST | 0.59 | 0.75 | 0.20 | 0.34 | 0.59 | 0.89 | 0.38 | 0.21 | 0.78 | 0.75 | 0.67 |
| THItoGene | 0.14 | 0.66 | 0.14 | 0.21 | 0.36 | 0.89 | 0.39 | 0.16 | 0.77 | 0.74 | 0.55 |
| MISO | 0.67 | 0.90 | 0.33 | 0.59 | 0.80 | 0.90 | 0.43 | 0.32 | 0.88 | 0.87 | 0.84 |
| HiST | 0.58 | 0.68 | 0.09 | 0.75 | 0.65 | 0.95 | 0.49 | 0.48 | 0.68 | 0.94 | 0.79 |
| sCellST | 0.89 | 0.89 | 0.45 | 0.42 | 0.87 | 0.87 | 0.38 | 0.22 | 0.87 | 0.78 | 0.82 |
| iStar | 0.81 | 0.83 | 0.33 | 0.51 | 0.80 | 0.95 | 0.41 | 0.41 | 0.88 | 0.92 | 0.86 |

**Perturbed H&E images**

| | iLISI | S-batch | KBET | G-Con | Batch overall | cLISI | S-label | K-NMI | scGraph | Bio. overall | Overall |
|---|---|---|---|---|---|---|---|---|---|---|---|
| SpaWeaver | 0.80 | 0.85 | 0.45 | 0.61 | 0.93 | 0.95 | 0.45 | 0.47 | 0.90 | 0.98 | 0.96 |
| DeepPT | 0.04 | 0.76 | 0.07 | 0.47 | 0.43 | 0.88 | 0.42 | 0.24 | 0.86 | 0.82 | 0.62 |
| SpatialEx+ | 0 | 0.46 | 0 | 0.38 | 0.26 | 0.92 | 0.43 | 0.32 | 0.84 | 0.86 | 0.56 |
| Hist2ST | 0 | 0.57 | 0.02 | 0.24 | 0.26 | 0.92 | 0.41 | 0.20 | 0.76 | 0.77 | 0.52 |
| THItoGene | 0 | 0.70 | 0.01 | 0.27 | 0.30 | 0.93 | 0.40 | 0.19 | 0.61 | 0.72 | 0.51 |
| MISO | 0 | 0.42 | 0 | 0.38 | 0.25 | 0.89 | 0.43 | 0.26 | 0.87 | 0.84 | 0.54 |
| HiST | 0.17 | 0.60 | 0.02 | 0.74 | 0.48 | 0.95 | 0.49 | 0.45 | 0.61 | 0.91 | 0.70 |
| sCellST | 0.86 | 0.86 | 0.37 | 0.44 | 0.86 | 0.87 | 0.39 | 0.22 | 0.88 | 0.79 | 0.82 |
| iStar | 0 | 0.69 | 0.02 | 0.47 | 0.37 | 0.95 | 0.44 | 0.34 | 0.87 | 0.90 | 0.63 |
